# β-catenin obstructs γδ T cell immunosurveillance in colon cancer through loss of BTNL expression

**DOI:** 10.1101/2022.06.10.495115

**Authors:** Toshiyasu Suzuki, Anna Kilbey, Rachel A. Ridgway, Hannah Hayman, Ryan Bryne, Nuria Casa Rodríguez, Anastasia Georgakopoulou, Lei Chen, Michael Verzi, David Gay, Ester G. Vázquez, Hayley L. Belnoue-Davis, Kathryn Gilroy, Anne Helene Køstner, Christian Kersten, Chanitra Thuwajit, Ditte Andersen, Robert Wiesheu, Anett Jandke, Natalie Roberts, Karen Blyth, Antonia Roseweir, Simon J. Leedham, Philip D. Dunne, Joanne Edwards, Adrian Hayday, Owen J. Sansom, Seth B. Coffelt

## Abstract

WNT/β-catenin signaling endows cancer cells with proliferative capacity and immune-evasive functions that impair anti-cancer immunosurveillance by conventional, cytoxtoic T cells. However, the impact of dysregulated WNT signalling on unconventional, tissue-resident T cells, specifically in colon cancer is unknown. Here, we show that cancer cells in *Apc*-mutant mouse models escape immunosurveillance from gut-resident intraepithelial lymphocytes (IELs) expressing γδ T cell receptors (γδTCRs). Analysis of late-stage tumors from mice and humans revealed that γδIELs are largely absent from the tumor microenvironment, and that butyrophilin-like (BTNL) molecules, which can critically regulate γδIEL through direct γδTCR-interactions, are also downregulated. We could attribute this to β-catenin stabilization, which rapidly decreased expression of the transcription factors, HNF4A and HNF4G, that we found to bind promoter regions of *Btnl* genes, thereby driving their expression in normal gut epithelial cells. Indeed, inhibition of β-catenin signaling restored *Btnl1* gene expression and γδ T cell infiltration into tumors. These observations highlight an immune-evasion mechanism specific to WNT-driven colon cancer cells that disrupts γδIEL immunosurveillance and furthers cancer progression.

## INTRODUCTION

The mammalian intestinal tract contains groups of tissue-resident T cells, called intraepithelial lymphocytes (IELs), which share a symbiotic relationship with the epithelial cell layer. IELs expressing the γδ T cell receptor (TCR) account for nearly 50% of all T cells in the mouse gut and 10-30% of all T cells in the human intestinal tract. These cells actively migrate in the space between the enterocyte layer and the basement membrane, surveying for abnormalities. γδIELs play instrumental roles in a multitude of physiological processes, such as homeostasis, epithelial cell shedding, infection, maintaining gut barrier integrity, nutrient sensing, dietary metabolism and tumor control (1–8).

Although diverse, the TCRs of most mouse γδIELs include a Vγ7 chain that facilitates critical interactions with butyrophilin-like (BTNL) molecules – specifically, heterodimers consisting of BTNL1 with BTNL4 or BTNL6 (9–11). Vγ7^+^ cells ordinarily reside only in gut tissue, owing at least in part to the largely restricted expression of BTNL1, BTNL4 and BTNL6 to intestinal epithelial cells (9–12). The BTNL1/6 or BTNL1/4 interaction drives Vγ7^+^ γδ IEL expansion and maturation during post-natal development and is thereafter required for maintaining the signature phenotype of Vγ7^+^ IEL (11, 13). The BTNL1/6-γδ T cell axis in mice is also conserved in humans: human BTNL3 and BTNL8 dimers bind to and regulate Vγ4^+^ IELs (11,12,14). The localization of γδ IELs and of BTNL expression aligns with a decreasing WNT signalling gradient that runs from crypt to villus. As such, Vγ7^+^ IELs are rarely found in the crypt regions where WNT signaling is high.

Most colorectal carcinomas exhibit mutations in members of the WNT pathway that drive tumor initiation and progression to malignancy. These mutations are almost exclusively manifest in the form of truncating mutations in the *APC* tumor suppressor gene, preventing the degradation of β-catenin, which leads to uncontrolled proliferation (15). Like intestinal stem cells residing in crypt regions, colon cancer cells require WNT signaling to maintain their stemness and de-differentiated phenotype (16, 17). Additionally, aberrant WNT signaling not only affects mutated epithelial cells, but it can also counteract immune surveillance and thwart anti-tumor immunity by dendritic cells and conventional CD8^+^ T cells in several cancer types (18–21). However, the relationship between dysregulated WNT signaling in cancer and local, tissue-resident IELs remains wholly unexplored.

Here, we investigated Vγ7^+^ IEL function and the expression of BTNL molecules during tumor initiation and growth. We found that β-catenin signaling in intestinal epithelial cells decreases expression of *Btnl* genes and the transcription factors that regulate them, HNF4A and HNF4G. This molecular rewiring promoted γδ T cell exclusion from tumors. Conversely, inhibition of β-catenin signaling restored HNF4 transcription factor expression, *Btnl1* gene expression and intra-tumoral γδ T cell infiltration. Collectively, our data suggest that aberrant WNT signaling in tumors elicits disarray in the tissue-resident γδ T cell compartment, disrupting natural tissue immunosurveillance as cancer cells dedifferentiate and acquire stem cell-like characteristics.

## RESULTS

### Vγ7^+^ cells suppress gut tumor formation

To test the importance of gut-resident Vγ7^+^ cells in tumor initiation and progression, we crossed *Villin-Cre^ERT2^;Apc^F/+^* (VA) mice with *Btnl1^—/—^*mice, which harbor significantly diminished Vγ7^+^ cell compartments in the small intestine (SI) and colon (11). Tumors were induced in VA and VA;*Btnl1^—/—^* mice by tamoxifen, and these mice were aged to humane endpoint. We confirmed that *Btnl1* expression is absent from gut tissue of VA;*Btnl1^—/—^* mice, while *Btnl1* expression is maintained in VA mice (Figure 1A). The number of γδ T cells in normal, tumor-adjacent regions was reduced in VA;*Btnl1^—/—^*mice when compared to VA mice (Figure 1B). Overall survival of tumor-bearing VA and VA;*Btnl1^—/—^* mice was the same, and there was comparable tumor incidence and burden in the SI of tumor-bearing VA and VA;*Btnl1^—/—^* mice (Figure 1C, D). Conversely, tumor number and particularly tumor burden were increased in the colon of VA;*Btnl1^—/—^* mice when compared to VA mice (Figure 1D). The lack of phenotype in the SI may be explained by compensation from cytotoxic TCRαβ^+^ IELs and other γδ T cell subsets (e.g. Vγ1^+^ cells), which partially offset Vγ7^+^ cell deficiencies in *Btnl1*-deficient mice (11). Since bacterial load is higher in the murine distal colon than in the SI (22), the propensity for inflammation-driven tumors in this anatomical location may be more sensitive to the lack of Vγ7^+^ cells, which are crucial infection sensors and protectors from pathogens (4). In sum, the BTNL1-Vγ7 axis evidently contributes to immunosurveillance during tumor initiation and growth.

**Figure 1.**
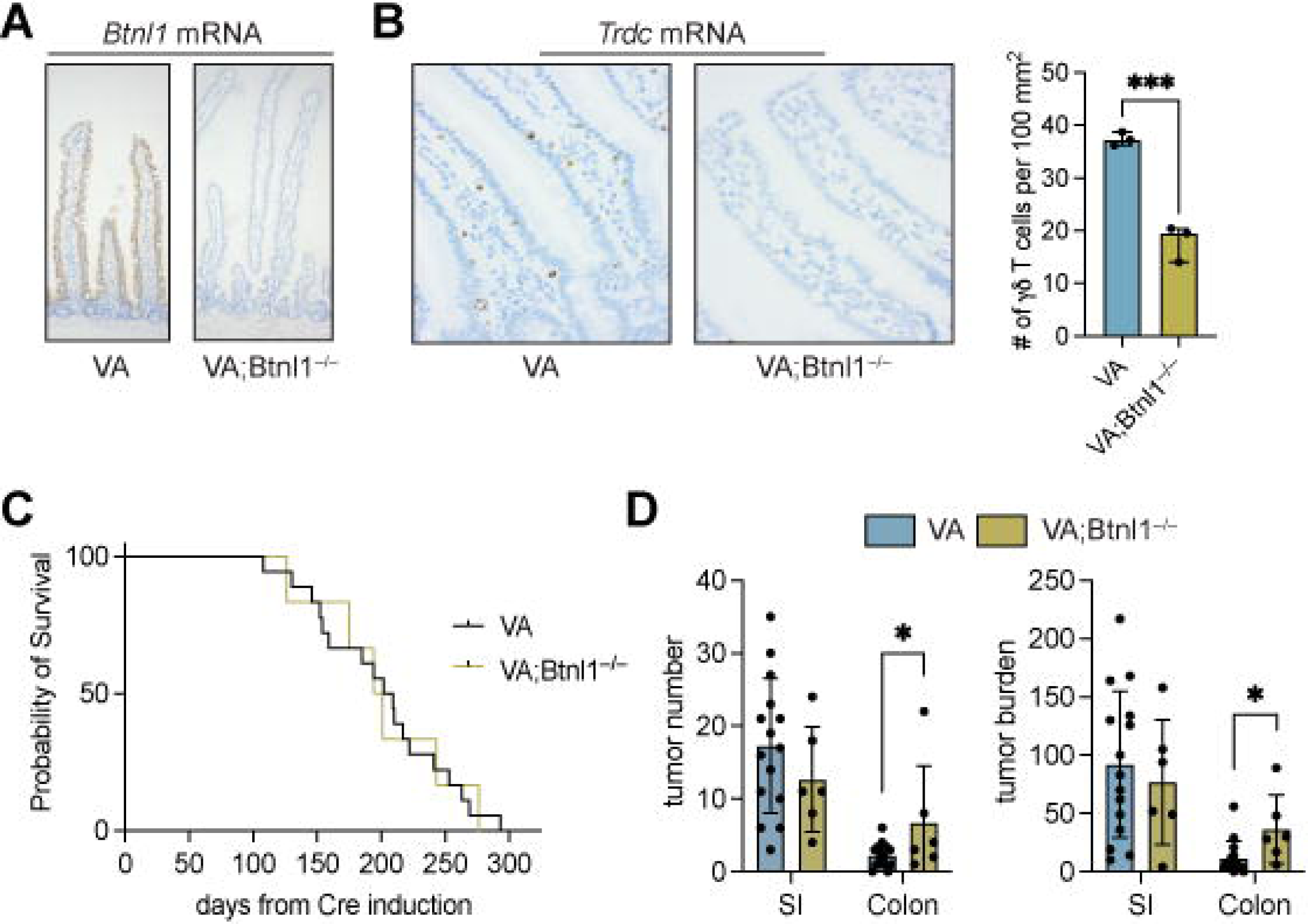
Loss of *Btnl1* increases adenoma formation in *Apc*-deficient mouse models. (A) Representative images of intestinal tissue from 4 VA and VA;*Btnl1^—/—^* mice stained for *Btnl1* mRNA. (B) Representative images of intestinal tissue from 4 VA and VA;*Btnl1^—/—^*mice stained for *Trdc* mRNA. Graphic representation of γδ T cell numbers in intestinal tissue of VA and VA;*Btnl1^—/—^* mice. Each dot represents one mouse (n = 3). Data presented as mean ± SD per 100 mm^2^. \*\*\**p* < 0.001 as determined by unpaired t test. (C) Kaplan-Meier survival analysis of VA and VA;*Btnl1^—/—^*mice (n = 15 VA, 6 VA;*Btnl1^—/—^* mice). (D) Graphic representation of tumor number and tumor burden in the small intestine (SI) and colon of VA and VA;*Btnl1^—/—^*mice. Each dot represents one mouse (n = 15 VA, 6 VA;*Btnl1^—/—^* mice). Data presented as mean ± SD. \**p* < 0.05 as determined by unpaired t test.

### Mouse and human tumors exhibit a paucity of γδ T cells

We next asked whether the prevalence of Vγ7^+^ cells in normal gut tissue was maintained in tumors. Contrary to their abundance in gut tissue of wild-type (WT) mice, γδ T cells were sparse within adenomas from VA mice, as well as an additional model of colon cancer, *Villin-Cre^ERT2^;Apc^F/+^;Kras^G12D/+^* (VAK) mice (Figure 2A). The cells’ frequency within tumors was estimated at 7-10 fold lower than in normal tissue (Figure 2B). Because Vγ7^+^ IELs express CD8αα dimers, whereas most other intestinal γδ T cells do not (11), we used CD8α as a marker to specifically quantify the Vγ7^+^ cell representation in tumor-bearing VA and VAK mice. CD8^+^ γδ T cells were apparent in the SI of WT or tumor-bearing VA and VAK mice; however, CD8^+^ γδ T cells were almost absent from tumors in either the VA or VAK model (Figure 2C). These observations show that tumor-infiltrating Vγ7^+^ cells and other γδ T cell subsets are rare.

**Figure 2.**
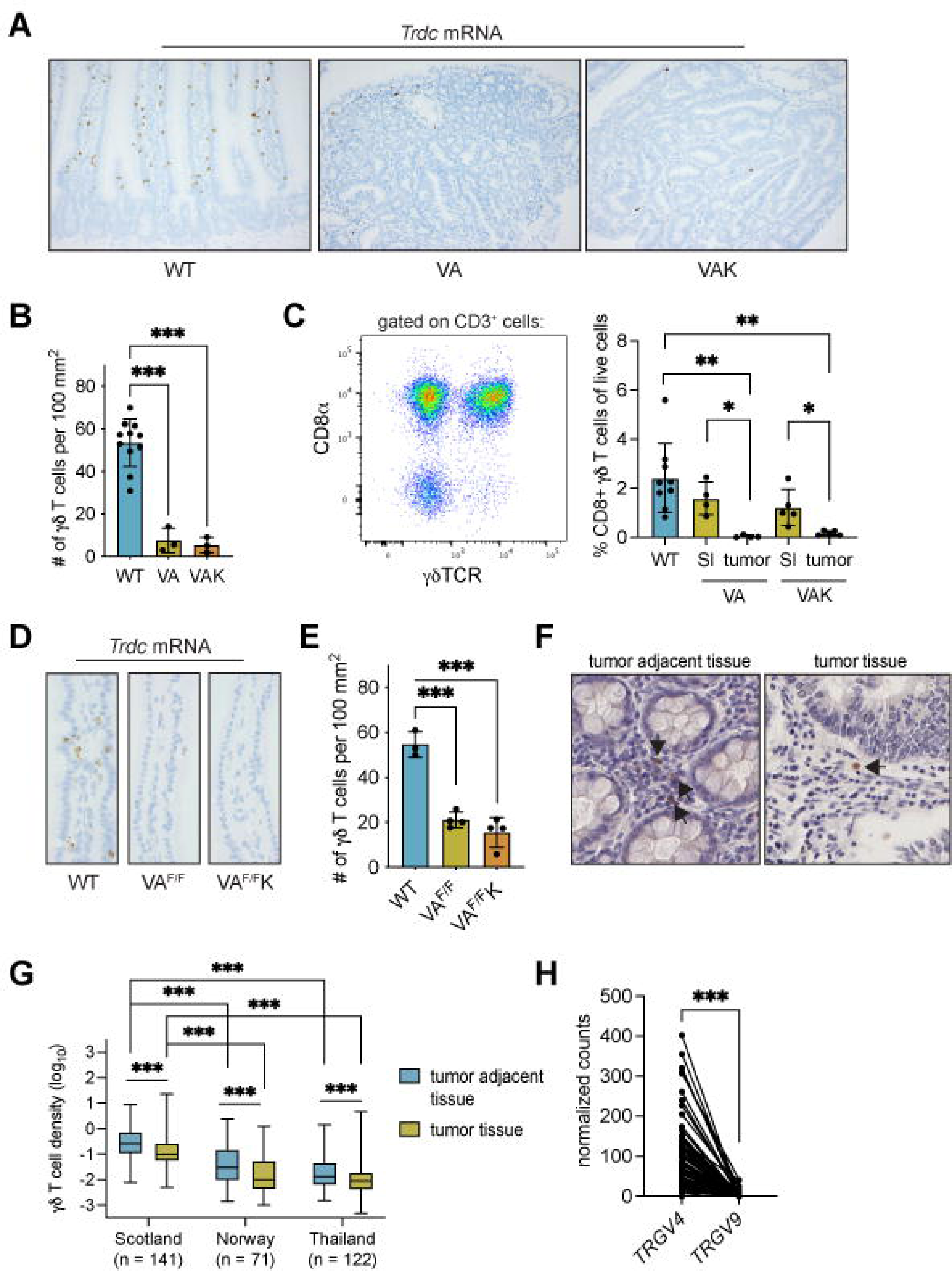
γδ T cells are excluded from mouse and human gut tumors. (A) Representative images of intestinal tissue from 4 wild-type (WT, Cre negative), tumor-bearing *Villin-Cre^ERT2^;Apc^F/+^*(VA) and tumor-bearing *Villin-Cre^ERT2^;Apc^F/+^;Kras^G12D^*(VAK) mice stained for *Trdc* mRNA. (B) Graphic representation of γδ T cell numbers in intestinal tissue of WT mice and in tumors of VA and VAK mice. Each dot represents one mouse (n = 11 WT, 3 VA, 3 VAK). Data presented as mean ± SD per 100 mm^2^. \*\*\**p* < 0.001 as determined by one-way ANOVA followed by Dunnett’s posthoc test. (C) Representative flow cytometry plot of CD8α and γδTCR expression on total CD3^+^ cells in the small intestine of WT mice. Frequency of γδ T cells expressing CD8α in the small intestine (SI) of WT mice, as well as the SI and tumor of VA and VAK mice. Each dot represents one mouse (n = 9 WT, 4 VA, 5 VAK). Data presented as mean ± SD. \**p* < 0.05, \*\**p* < 0.01 as determined by one-way ANOVA followed by Dunnett’s posthoc test. (D) Representative images of SI from 4 wild-type (WT), *Villin-Cre^ERT2^;Apc^F/F^*(VA^F/F^) and *Villin-Cre^ERT2^;Apc^F/F^;Kras^G12D^* (VA^F/F^K) mice stained for *Trdc* mRNA. (E) Graphic representation of γδ T cell numbers in intestinal tissue of WT, VA^F/F^ and VA^F/F^K mice. Each dot represents one mouse (n = 3 WT, 4 VA^F/F^, 4 VA^F/F^K). Data presented as mean ± SD per 100 mm^2^. \*\*\**p* < 0.001 as determined by one-way ANOVA followed by Dunnett’s posthoc test. (F) Representative image of γδ T cell staining in tumor adjacent tissue and tumor tissue from 141 human colon cancer sections where arrows indicate positively stained cells. (G) Density of γδ T cells in human colon cancer sections in three different patient cohorts: Scotland (n = 141), Norway (n = 71) and Thailand (n = 122). γδ T cells identified by IHC in full sections were quantified in tumor adjacent tissue or tumor tissue using Visiopharm. Data presented as median ± min/max. \*\*\**p* < 0.001 as determined by paired t test. (H) Expression of *TRGV4* and *TRGV9* mRNA in human colon cancer samples (n = 82) from the Scotland cohort determined by TempO-Seq. \*\*\**p* < 0.001 as determined by paired t test.

How quickly γδ T cells might be excluded from tumors was investigated using a short-term model, wherein both alleles of *Apc* are simultaneously deleted in gut tissue, thereby maximally activating β-catenin signaling. The mouse intestine cannot tolerate loss of *Apc* in this way, so mice are culled 3 or 4 days after CRE recombinase induction. γδ T cells were quantified in villi of the SI of *Villin-Cre^ERT2^;Apc^F/F^*(VA^F/F^) mice and *Villin-Cre^ERT2^;Apc^F/F^;Kras^G12D/+^*(VA^F/F^K) mice. The number of γδ T cells was reduced by about 3-fold in VA^F/F^ and VA^F/F^K mice when compared to WT controls (Figure 2D, E), indicating that deletion of *Apc* in epithelial cells has a rapid impact on γδ T cell numbers, prior to the overt formation of a tumor.

To investigate whether our findings might find parallels in human colon tumors, we examined samples from three human cohorts that were collected from Scotland, Norway and Thailand. γδ T cells were quantified in tumor tissue and normal adjacent tissue after immunohistochemistry with a pan-γδ T cell antibody using digital pathology software (Figure 2F). In all three cohorts, γδ T cell densities were higher in normal adjacent tissue than tumor tissue (Figure 2G), mirroring our observations in mouse models. Moreover, levels of γδ T cells were higher in the Scotland cohort when compared to Norway and Thailand cohorts (Figure 2G). We performed RNAseq analysis on 82 human colon cancer samples from the Scotland cohort from which immunohistochemical γδ T cell density data were available to glean information on the subtypes of γδ T cells present in these tumors. *TRGV4* transcripts were more abundant than *TRGV9* transcripts within the same tumor (Figure 2H), indicating that Vγ4^+^Vδ1^+^ cells, which reflect colonic IEL, are on aggregate more abundant than Vγ9^+^Vδ2^+^ cells which are typical of peripheral blood. These data corroborate but substantially extend findings by others (23, 24).

### *Btnl* molecules are downregulated in colorectal cancer

With a paucity of γδ T cells in the intestinal tumor microenvironment conserved across mouse and humans, we next investigated the expression of *Btnl* genes that are essential to the phenotypic maintenance of Vγ7^+^ IEL the adult gut (10,11,13). When tumor sections from VA and VAK mice were stained for *Btnl1* mRNA, expression was apparent in epithelial cells surrounding adenomas but was absent from cancer cells in both VA and VAK models (Figure 3A). This lack of *Btnl1* expression in tumors can be viewed as a contributory factor to the dysregulation of the CD8αα^+^Vγ7^+^ IEL compartment in the tumor microenvironment.

**Figure 3.**
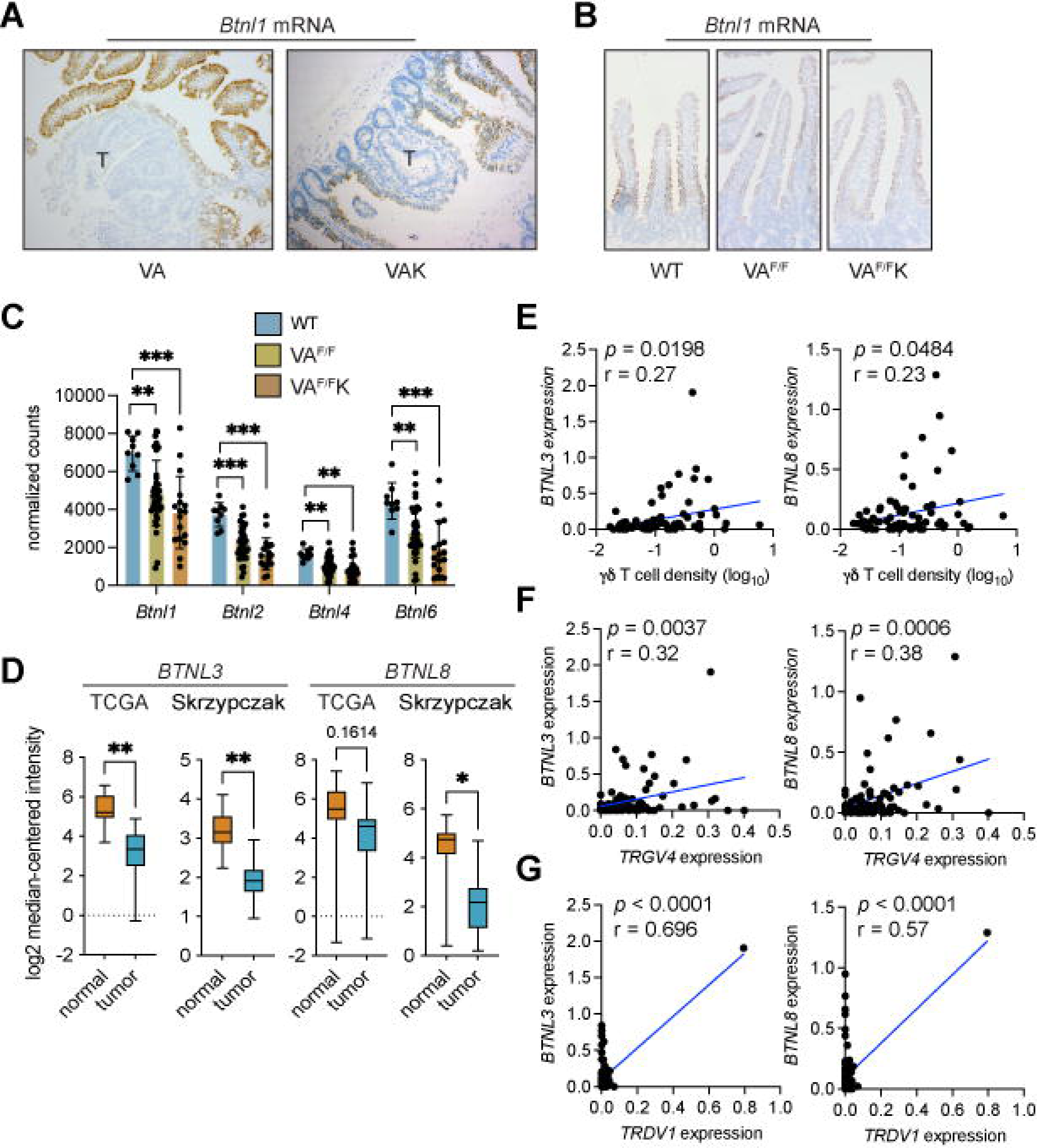
Expression of butyrophilin-like molecules is reduced in gut tumors. (A) Representative images of intestinal tissue from 4 VA and VAK mice stained for *Btnl1* mRNA. T = tumor. (B) Representative images of intestinal tissue from 4 WT, VA^F/F^ and VA^F/F^K mice stained for *Btnl1* mRNA. (C) Butyrophilin-like mRNA expression shown by heatmap generated from RNAseq data from WT, VA^F/F^ and VA^F/F^K mice (n = 3 mice/group). (D) Expression of *BTNL3* and *BTNL8* in normal human colonic tissue and tumor tissue from TCGA (n = 19 normal, 101 tumor) and Skrypczak (n = 24 normal, 45 tumor) datasets. Data presented as median ± min/max. \**p* < 0.05, \*\**p* < 0.01 as determined by Mann-Whitney U test. (E) Correlation between *BTNL3* or *BTNL8* expression as determined by TempO-Seq and γδ T cell density determined by IHC in the Scotland cohort from 77 matched pairs. Units on axes are normalized read counts x 10^3^. Each dot represents one tumor. *P* value and r value determined by Pearson’s correlation. (F) Correlation between *BTNL3* or *BTNL8* expression and *TRGV4* expression as determined by TempO-Seq in the Scotland cohort. Units on axes are normalized counts x 10^3^. Each dot represents one tumor (n = 82). *P* value and r value determined by Pearson’s correlation. (G) Correlation between *BTNL3* or *BTNL8* expression and *TRDV1* expression as determined by TempO-Seq in the Scotland cohort. Units on axes are normalized counts x 10^3^. Each dot represents one tumor (n = 82). *P* value and r value determined by Pearson’s correlation.

The kinetics of this loss in *Btnl1* expression were examined in the short-term VA^F/F^ and VA^F/F^K models. In these models, deletion of two copies of *Apc* with or without expression of mutant KRAS resulted in a slight reduction of *Btnl1* expression (Figure 3B). To verify this reduction, gene expression of *Btnl1* and of *Btnl2, Btnl4* and *Btnl6* were analyzed in RNAseq data from the SI of WT, VA^F/F^ and VA^F/F^K mice (25, 26). This analysis showed reduced RNA expression of all four *Btnl* family members following deletion of *Apc* in gut tissue (Figure 3C).

We next interrogated two human gene expression datasets (15, 27) to determine whether *BTNL3* or *BTNL8* – homologs of mouse *Btnl1* and *Btnl6* – expression levels were different between normal gut tissue and tumor tissue. *BTNL3* expression levels were higher in normal tissue than tumor tissue in both the TCGA and Skrzypczak datasets, while *BTNL8* expression was only higher in normal tissue in the Skrzypczak dataset (Figure 3D). These findings are similar to observations made by others (28). Together, our analyses demonstrate an evolutionarily conserved reduction of *BTNL* expression in tumors across species.

The relationship between expression of *BTNL3* and *BTNL8* and γδ T cell infiltration into human tumors was investigated in the Scotland cohort. Gene expression values from 77 human colon cancer samples were plotted with γδ T cell density values from matched samples. Both *BTNL3* and *BTNL8* mRNAs were positively correlated with γδ T cell density, with human tumors exhibiting high expression of *BTNL3* and *BTNL8* containing more γδ T cells than tumors with low levels of *BTNL3* and *BTNL8* (Figure 3E). To more specifically address the relationship between Vγ4^+^Vδ1^+^ IELs, *BTNL3* and *BTNL8* levels, we compared *TRGV4* and *TRDV1* expression levels with *BTNL3/8* expression levels. *TRGV4* mRNA was positively correlated with both *BTNL3* and *BTNL8* expression (Figure 3F). *TRDV1* mRNA was also positively correlated with both *BTNL3* and *BTNL8* expression; although, *TRDV1* mRNA was not detected in 33 of 82 samples (Figure 3G). These data support the notion that loss of *BTNL* molecules in tumors is directly associated with the loss of Vγ4^+^Vδ1^+^ IELs in the tumor microenvironment of human tumors.

### β-catenin signaling negatively regulates *Btnl* expression

The data reported above suggested a relationship between WNT signaling and loss of BTNL molecules in cancer cells with γδ T cell exclusion from the tumors, since loss of *Apc* resulted in rapid reduction of these molecules and cells. To explore this relationship in greater detail, we asked whether there was a correlation between the WNT pathway and γδ T cell density in human tumors. Expression levels of *CTNNB1* (which encodes β-catenin) and *SOX9* – a transcriptional target of the β-catenin transcription factor complex (29) – were plotted together with γδ T cell density values determined by immunohistochemistry from the same, matched tumor samples from the Scotland cohort. This analysis revealed that higher expression levels of *CTNNB1* and *SOX9* were correlated with low numbers of γδ T cells in human colon cancer (Figure 4A). Similarly, *TRGV4* mRNA negatively correlated with both *CTNNB1* and *SOX9* expression; although, this correlation did not reach significance for the *SOX9* comparison (Figure 4A). We did not explore correlations with *TRDV1* mRNA owing to the absence of detectable expression levels in many samples. In these human tumors, high *CTNNB1* and *SOX9* expression levels were correlated with low *BTNL3* and *BTNL8* expression (Figure 4B). To validate these findings, we analyzed the Marisa cohort, a publicly available gene expression dataset containing 258 human tumor samples (30). Within this dataset, high *CTNNB1* and *SOX9* expression levels correlated with low *BTNL3* expression levels, while *SOX9* also negatively correlated with *BTNL8* expression (Figure 4C). These data support our hypothesis of a relationship between WNT signalling, Vγ4^+^Vδ1^+^ IEL exclusion from tumors, and a loss of *BTNL* genes in cancer cells.

**Figure 4.**
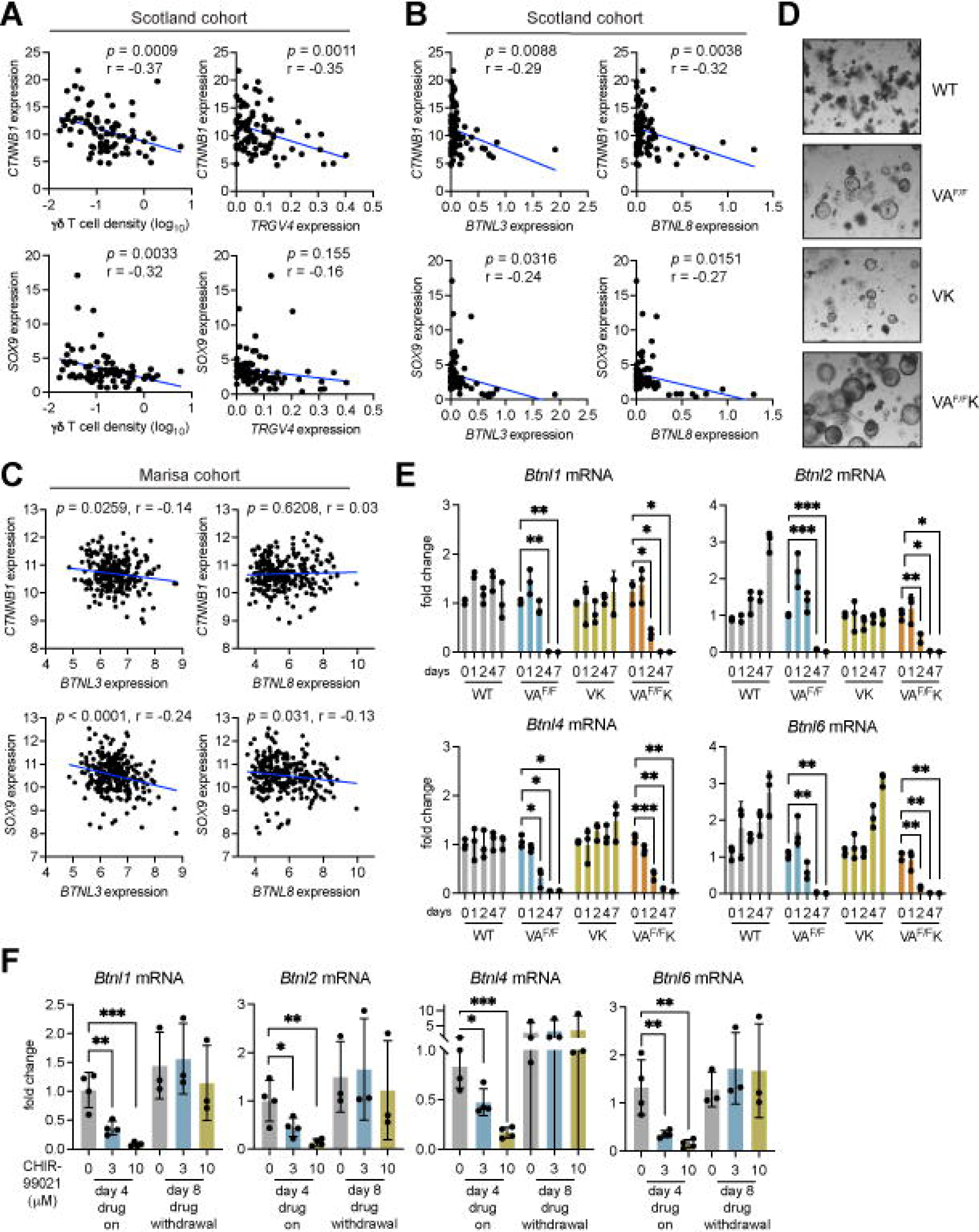
Activation of β-catenin decreases butyrophilin-like molecule expression. (A) Correlation between *CTNNB1* or *SOX9* and γδ T cell density or *TRGV4* expression as determined by TempO-seq and γδ T cell density determined by IHC in the Scotland cohort. Units on axes are normalized read counts x 10^3^. Each dot represents one tumor (n = 77 left panels, 82 right panels). *P* value and r value determined by Pearson’s correlation. (B) Correlation between *CTNNB1* or *SOX9* expression and *BTNL3* or *BTNL8* expression as determined by TempO-Seq in the Scotland cohort. Units on axes are normalized counts x 10^3^. Each dot represents one tumor (n = 82). *P* value and r value determined by Pearson’s correlation. (C) Correlation between *CTNNB1* or *SOX9* expression and *BTNL3* or *BTNL8* expression in the Marisa cohort. Units on axes are normalized counts x 10^3^. Each dot represents one tumor (n = 258). *P* value and r value determined by Pearson’s correlation. (D) Representative images of organoids derived from WT, VA^F/F^, VK and VA^F/F^K mice. Images were taken 4 days after tamoxifen treatment. (E) Fold change in expression levels of indicated genes in WT, VA^F/F^, VK and VA^F/F^K organoids. Gene expression was measured at indicated days post tamoxifen treatment. Each dot represents one organoid derived from one mouse. Data presented as mean ± SD. \**p* < 0.05, \*\**p* < 0.01 and \*\*\**p* < 0.001 as determined by one-way ANOVA followed by Dunnett’s posthoc test. (F) Fold change in expression levels of indicated genes in WT organoids treated with 3 or 10 μM CHIR-99021 for indicated days. Each dot represents one organoid derived from one mouse. Data presented as mean ± SD. \**p* < 0.05, \*\**p* < 0.01 as determined by one-way ANOVA followed by Dunnett’s posthoc test.

To explore a mechanistic link between WNT signalling activation and the down-regulation of *Btnl* gene expression, we developed an *ex vivo* transformation assay using intestinal organoids derived from tamoxifen-naïve WT, VA^F/F^, VK and VA^F/F^K mice. Cells were treated with tamoxifen *in vitro* to induce deletion of *Apc* or expression of mutant KRAS via Cre recombinase. Tamoxifen treatment failed to influence the shape or size of organoids derived from WT mice (Figure 4D). By contrast, tamoxifen altered the morphology of organoids harboring transgenic alleles, transforming their normal, budding shape into large spheres typical of tumor-derived organoids (Figure 4D). Gene expression was measured in these four groups of organoids over the course of one week after tamoxifen treatment. We confirmed that WNT pathway target genes, including *Lgr5, Sox9, Axin2* and *Cd44,* were up-regulated in VA^F/F^ and VA^F/F^K organoids, without affecting the same genes in WT and VK organoids (Supplemental Figure 1A). These results show that the organoid system recapitulates cancer cell transformation *in vivo* by β-catenin signaling. Expression of *Btnl1, Btnl2, Btnl4* and *Btnl6* mRNA was measured in these four groups of organoids (Figure 4E). Whereas expression of these genes remained constant in WT organoids, the deletion of *Apc* resulted in reduced expression of all *Btnl* RNAs assayed by day 4. Interestingly, activation of mutant KRAS had no effect on *Btnl* expression. However, the combination of *Apc* deletion and mutant KRAS expression in VA^F/F^K organoids accelerated *Btnl* down-regulation with reduced expression apparent by day 2 (Figure 4E). These observations demonstrate that β-catenin activation *via* loss of *Apc* negatively regulates *Btnl* gene expression.

As an alternative approach to genetic manipulation of WNT signaling, organoids from WT mice were treated with the GSK3β inhibitor, CHIR-99021, to activate β-catenin. Expression of *Btnl1, Btnl2, Btnl4* and *Btnl6* mRNA was measured after four days of treatment using two different concentrations of CHIR-99021. Both concentrations reduced expression of all *Btnl* RNAs assayed when compared to controls (Figure 4F), thus supporting the hypothesis that activated β-catenin down-regulates *Btnl gene* expression. The reversibility of this effect was tested by treating WT organoids with CHIR-99021 for 4 days, washing off drug, then culturing the treated organoids for another 4 days without drug. On day 8 after treatment began, expression of *Btnl1, Btnl2, Btnl4* and *Btnl6* mRNA was measured by qPCR. Withdrawal of CHIR-99021 restored *Btnl1, Btnl2, Btnl4* and *Btnl6* mRNA expression to baseline or higher levels in these organoids (Figure 4F).

### *Btnl* genes are regulated by HNF4 transcription factors

To understand how WNT signaling negatively affects *Btnl* gene expression, we investigated how *Btnl* molecules are regulated in normal tissue. Mouse BTNL1, BTNL2, BTNL4 and BTNL6 and human BTNL3 and BTNL8 are expressed by enterocytes and colonocytes in the intestinal tract (10,11,13,31). We hypothesized that restriction of BTNL molecule expression to the gut is a consequence of regulation by gut-specific transcription factors. To understand which transcription factors are important for induction of *BTNL* genes, we searched for potential transcription factor binding sites in the promoter regions of these genes. Using a publicly available database (OregAnno), we generated a list a putative transcription factor binding sites, then narrowed down the list by focusing on gut-specific transcription factors. This analysis uncovered two sets of paralogs: CDX1 and CDX2; HNF4A and HNF4G. Multiple binding sites for these proteins were found within 12 kb upstream of mouse and human *BTNL* gene start sites (Figure 5A).

**Figure 5.**
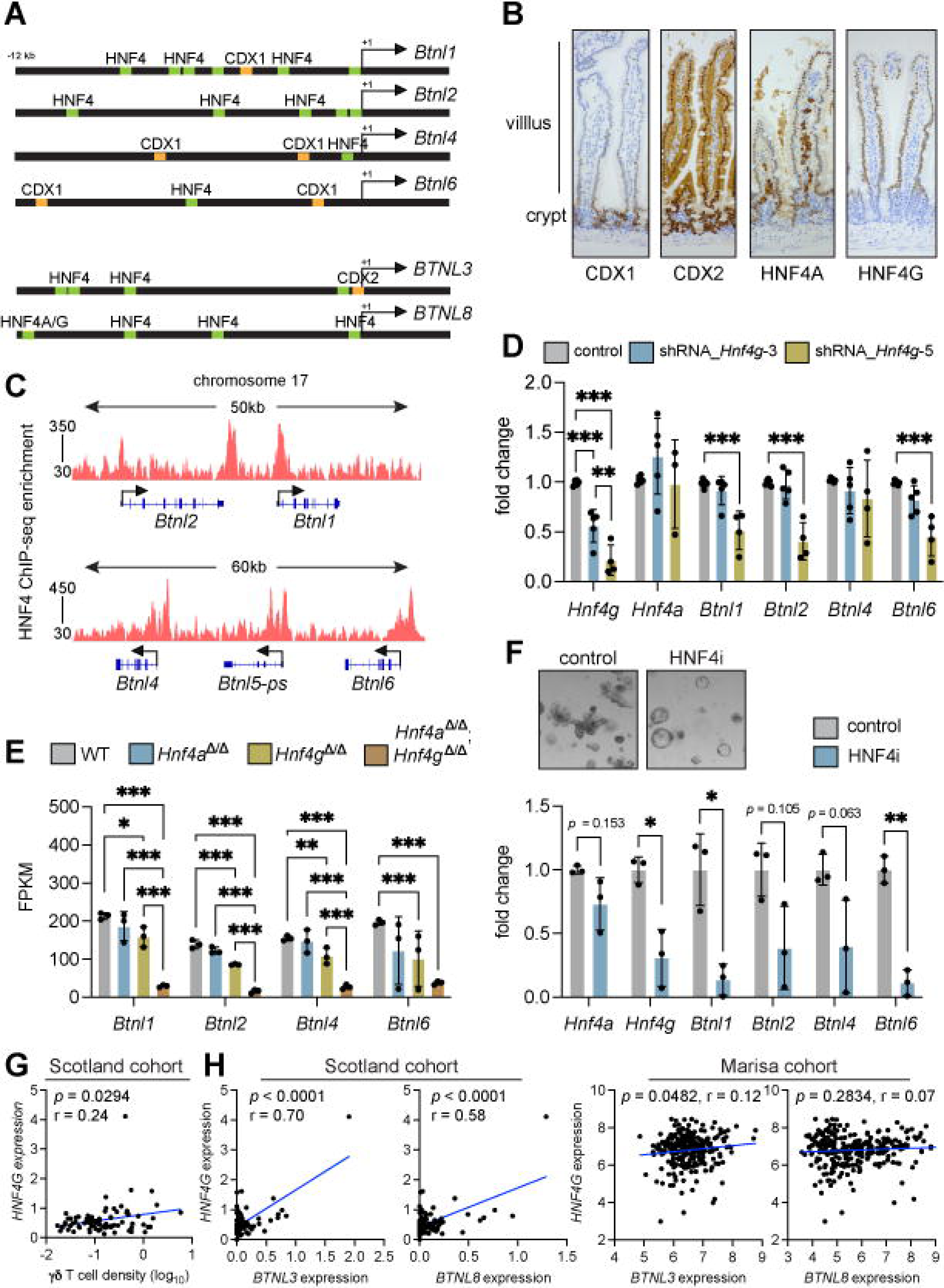
HNF4A and HNF4G regulate butyrophilin-like molecule expression in normal gut tissue. (A) Schematic of *Btnl1, Btnl2, Btnl4, Btnl6, BTNL3* and *BTNL8* promoter regions. Putative HNF4A/G binding sites are shown in green; CDX1 and CDX2 binding sites are shown in orange. (B) Representative images of CDX1, CDX2, HNF4A and HNF4G protein expression in small intestine of 4 WT mice. (C) Integrative Genomics Viewer analysis of HNF4A/HNF4G ChIP-seq data at mouse *Btnl* gene loci. (D) Fold change in expression levels of indicated genes in WT organoids transduced with shRNA constructs targeting *Hnf4g* transcripts. Each dot represents one organoid from one mouse. Data presented as mean ± SD. \*\**p* < 0.01, \*\*\**p* < 0.001 as determined by one-way ANOVA followed by Tukey’s posthoc test. (E) Butyrophilin-like molecule expression determined by RNAseq analysis of small intestine in WT, *Villin-Cre^ERT2^;Hnf4a^F/F^*(*Hnf4a*^Δ*/*Δ^), *Hnf4g^Crispr/Crispr^*(*Hnf4g*^Δ*/*Δ^), *Hnf4a*^Δ*/*Δ^*;Hnf4g*^Δ*/*Δ^ mice. Each dot represents one mouse. Data presented as mean ± SD. \**p* < 0.05, \*\*\**p* < 0.001 as determined by one-way ANOVA followed by Tukey’s posthoc test. (F) Representative images of organoids from WT mice treated with DMSO control or HNF4A/G inhibitor (HNF4i). Fold change in expression levels of indicated genes Each dot represents one organoid from one mouse. Data presented as mean ± SD. * < 0.05, \*\**p* < 0.01 as determined by unpaired t test. (G) Correlation between *HNF4G* expression as determined by TempO-seq and γδ T cell density determined by IHC in the Scotland cohort. Units on axes are normalized counts x 10^3^. Each dot represents one tumor (n = 77). *P* value and r value determined by Pearson’s correlation. (H) Correlation between *BTNL3* or *BTNL8* expression and *HNF4G* expression Units on axes are normalized counts x 10^3^. Each dot represents one tumor (n = 82 Scotland cohort, 258 Marisa cohort). *P* value and r value determined by Pearson’s correlation.

BTNL molecules are expressed in differentiated regions of the villus, not in the stem cell regions of the crypt. Therefore, we determined whether any or all of CDX1, CDX2, HNF4A or HNF4G were localized specifically to the villus. The protein expression pattern of each molecule was investigated in mouse intestine. CDX1 was expressed in crypt regions and lower villus, but expression decreased as enterocytes moved up the villus (Figure 5B). CDX2 was expressed in both crypts and villi with higher expression in the crypt. HNF4A was also expressed in both crypts and villi; staining was also observed in cells residing within the lamina propria. HNF4G expression was specific to enterocytes in the villus, as no expression was observed in crypt regions (Figure 5B). These data suggest that HNF4G is the prime candidate for *Btnl* gene regulation given their overlapping patterns of expression in the villus. However, all four transcription factors are expressed in the villus to some extent.

We investigated whether CDX1 and CDX2 mediate *Btnl1, Btnl2, Btnl4* and *Btnl6* transcription. Organoids from WT mice were transduced with 5 shRNA constructs targeting *Cdx1* or *Cdx2* mRNA. Two constructs achieved good knockdown efficiency for *Cdx1*; although, organoid morphology and expression of *Btnl* molecules remained unchanged (Supplemental Figure 2A, B). Attempts to knockdown *Cdx2* proved difficult as organoids transduced with these constructs often died. In two replicate experiments where organoids survived antibiotic selection, knockdown of *Cdx2* was sufficiently achieved with the shRNA_*Cdx2*-2 construct, but this failed to impact on organoid morphology or *Btnl* gene expression (Supplemental Figure 2C, D). These data suggest that CDX2 is required for organoid survival. Indeed, conditional deletion of *Cdx2* in adult intestine is lethal (32). We concluded from these experiments that CDX1 and CDX2 are not required specifically for *Btnl1, Btnl2, Btnl4* and *Btnl6* transcription.

HNF4A and HNF4G are paralogs that bind fatty acids and whose functions are somewhat redundant (33–35). These transcription factors recognize a nearly identical consensus motif on DNA, and they exhibit 98.7% commonality in DNA binding profiles (33). It should be noted that HNF4A is expressed outside the gut at sites such as liver (36), where BTNL molecules are not expressed (10). To determine whether HNF4A and HNF4G bind the promoter region of *Btnl1, Btnl2, Btnl4* and *Btnl6* genes, we analyzed chromosome 17 in a HNF4 ChIP-seq dataset from mouse SI (33). This analysis confirmed that HNF4A/G bind all *Btnl* gene promoter regions (Figure 5C).

Having established that HNF4A/G occupy *Btnl* promoters, we investigated whether HNF4A and HNF4G activity was causally linked to *Btnl* expression. Organoids from WT mice were transduced with 5 shRNA constructs targeting *Hnf4a* or *Hnf4g* mRNA. Organoid morphology was unaffected by *Hnf4a* constructs (Supplemental 2E). Knockdown of *Hnf4a* was not successful. Instead of reduced expression, we observed higher expression of *Hnf4a* and *Hnf4g* in these cells, concomitant with higher expression of *Btnl1, Btnl2, Btnl4* and *Btnl6* genes (Supplemental 2F). These findings suggest that a feedback mechanism may be active, preventing *Hnf4a* knockdown, but provide indirect evidence that increased HNF4A and HNF4G expression correlates with increased *Btnl* expression. To clarify this situation, we transduced MODE-K enterocytes that do not express *Hnf4a* with a series of cDNAs encoding gut-associated transcription factors, including *Cdx1*, *Cdx2* and *Hnf4a*. Only in *Hnf4a* transductants was there overt upregulation of *Btnl* mRNAs, specifically those for *Btnl4* and *Btnl6* (Supplemental Figure 2G), while *Cdx1, Cdx2, Creb3l3, Gata5* and *Isx* failed to influence *Btnl* gene expression. For *Hnf4g* targeting in organoids from WT mice, two constructs achieved good knockdown efficiency with the shRNA_*Hnf4g-5* construct exhibiting superior efficiency. However, only the shRNA_*Hnf4g-5* construct reduced expression of *Btnl1*, *Btnl2* and *Btnl6* without affecting expression of *Btnl4* (Figure 5D). This was accompanied by the occasional appearance of sphere-shaped organoids (Supplemental Figure 2H), indicative of a stem cell-like state. Collectively, our data demonstrate that HNF4 transcription factors are regulators of *Btnl* gene expression; although, there are seemingly differences in the degrees to which specific *Btnl* genes are dependent upon or influenced by HNF4A and HNF4G, respectively. These data further integrate *Btnl* expression with physiologic enterocyte differentiation (33, 37).

We analyzed *Btnl* gene expression in mouse models deficient for HNF4A or HNF4G or both. An RNA-seq dataset derived from intestine of WT, *Villin-Cre^ERT2^;Hnf4a^F/F^*(*Hnf4a*^Δ*/*Δ^) mice, *Hnf4g^—/—^* (*Hnf4g*^Δ*/*Δ^) mice and *Hnf4a*^Δ*/*Δ^*;Hnf4g*^Δ*/*Δ^ mice was used for this purpose (33). In these mice, deletion of *Hnf4a* failed to alter *Btnl* gene expression, while deletion of *Hnf4g* reduced all four *Btnl* genes (Figure 5E). Simultaneous deletion of *Hnf4a* and *Hnf4g* led to the most pronounced loss of *Btnl* expression when compared to WT tissue. *Btnl1, Btnl2* and *Btnl4* (but not *Btnl6*) mRNA was also lower in *Hnf4a*^Δ*/*Δ^*;Hnf4g*^Δ*/*Δ^ intestine than in *Hnf4a*^Δ*/*Δ^ or *Hnf4g*^Δ*/*Δ^ intestine (Figure 5E). We then used an inhibitor that targets both HNF4A and HNF4G, BI-6015 (HNF4i), in organoids from WT mice. This drug altered the morphology of the organoids, transforming them into spheres (Figure 5F), similar to the morphology of *Hnf4a*^Δ*/*Δ^*;Hnf4g*^Δ*/*Δ^ organoids previously described (33). Inhibition of these transcription factors by BI-6015 reduced expression of *Hnf4a* and *Hnf4g*, as well as *Btnl1, Btnl2, Btnl4* and *Btnl6* mRNA (Figure 5F). Together, these data demonstrate that HNF4G is the main regulator of *Btnl* molecule expression with cooperation from HNF4A in enterocytes.

The relationship between transcription factor expression, γδ T cell infiltration and *BTNL* gene expression was examined in human tumors. In the Scotland cohort, there was no correlation between *CDX1, CDX2* and *HNF4A* expression and γδ T cell density (Supplemental Figure 3A-C). There was a positive correlation between *CDX1* expression, *BTNL3* expression and *BTNL8* expression in both the Scotland and Marisa cohorts (Supplemental Figure 3D), but correlations between *CDX2* or *HNF4A* and *BTNL3* or *BTNL8* were absent or inconsistent among both cohorts (Supplemental Figure 3E-F). In contrast to the other transcription factors, increased *HNF4G* expression was correlated with higher γδ T cell density in human tumors (Figure 5G). Increased *HNF4G* expression also correlated with increased *BTNL3* expression in both the Scotland and Marisa cohorts, whereas the relationship with *BTNL8* expression was only observed in the Scotland cohort (Figure 5H). These data establish a strong association between *HNF4G*, *BTNL* expression, and tumor-infiltrating γδ T cells in human tumors and point to HNF4G regulation of *BTNL* gene expression as being conserved across species.

### Increased WNT signaling disrupts HNF4 expression, *Btnl* expression and γδIELs

Having established how expression of *Btnl* molecules is controlled in differentiated regions of normal gut by HNF4A and HNF4G transcription factors, we hypothesized that disruption of the WNT gradient in the mouse intestine would interfere with enterocyte-specific HNF4G and *Btnl* gene expression and subsequently γδIEL abundance. To test this hypothesis, we used *Vil1-Grem1* mice in which *Gremlin1* is under the control of the *Villin* promoter. These mice develop ectopic crypts in the villi due to the antagonist actions of GREM1 on bone morphogenic proteins (BMPs), a consequence of which is increased WNT signaling in villi (38). Nuclear SOX9 was used to identify ectopic crypts in the villi of *Vil1-Grem1* mice (Supplemental Figure 4A). These SOX9-high, WNT-high ectopic crypts maintained HNF4A as in normal crypts, but lost expression of HNF4G and *Btnl1* mRNA (Supplemental Figure 4A). Moreover, we quantified γδ T cells in villi of *Vil1-Grem1* mice and found that these cells were reduced when compared to WT mice (Supplemental Figure 4A, B). These results show that WNT signaling suppresses the HNF4G-*Btnl*-Vγ7^+^ cell axis.

Further testing of our hypothesis was carried out in an additional model that is WNT ligand dependent in which R-spondin 3 (RSPO3) is expressed from LGR5^+^ stem cells: *Lgr5-Cre^ERT2^;Rspo3^INV^* mice (39). In this model, increased WNT signaling induces greater numbers of crypt regions, as demonstrated by increased SOX9^+^ cells at the base of the intestine, and reduced villus length (Supplemental Figure 4C). We investigated HNF4A, HNF4G and *Btnl1* expression in these mice. Staining patterns of these molecules were consistent with expression in intestine from WT mice where HNF4A was expressed in crypt regions and enterocytes, while HNF4G and *Btnl1* expression was restricted to enterocytes (Supplemental Figure 4C). However, the expansion of WNT-high crypt regions and reduced villus length resulted in fewer γδ T cells in villi of these in *Lgr5-Cre^ERT2^;Rspo3^INV^*mice, when compared to WT mice (4C, D). To determine whether reduced γδ T cell numbers could be restored by interference with over-expressed WNT ligands, *Lgr5-Cre^ERT2^;Rspo3^INV^* mice were treated with the porcupine inhibitor (PCPNi), LGK-974, to block the secretion of RSPO3 and prevent its activation of β-catenin. Expression patterns of HNF4A in crypt and villi regions as well as HNF4G and *Btnl1* mRNA in enterocytes were unaltered by LGK-974 treatment. However, γδ T cell numbers in the villi of these mice increased (Supplemental Figure 4C, D). These data provide evidence that aberrant β-catenin activation in normal intestinal tissue disrupts γδIEL abundance.

### *Hnf4a* and *Hnf4g* are suppressed by WNT signaling during tumor initiation

Given the similarities between crypt regions and gut tumors where WNT levels and β-catenin activity is high, we determined whether HNF4A and HNF4G are dysregulated in tumors. We hypothesized that β-catenin-induced loss of *Btnl* gene expression in tumors is a consequence of reduced HNF4A/HNF4G activity. To address this hypothesis, we initially compared *HNF4A* and *HNF4G* expression between normal human colon tissue and tumor tissue in the TCGA and Skrzypczak datasets. Both *HNF4A* and *HNF4G* were reduced in tumor tissue from both datasets (Figure 6A), mirroring reduced *BTNL3* and *BTNL8* expression in human tumors (Figure 3D).

**Figure 6.**
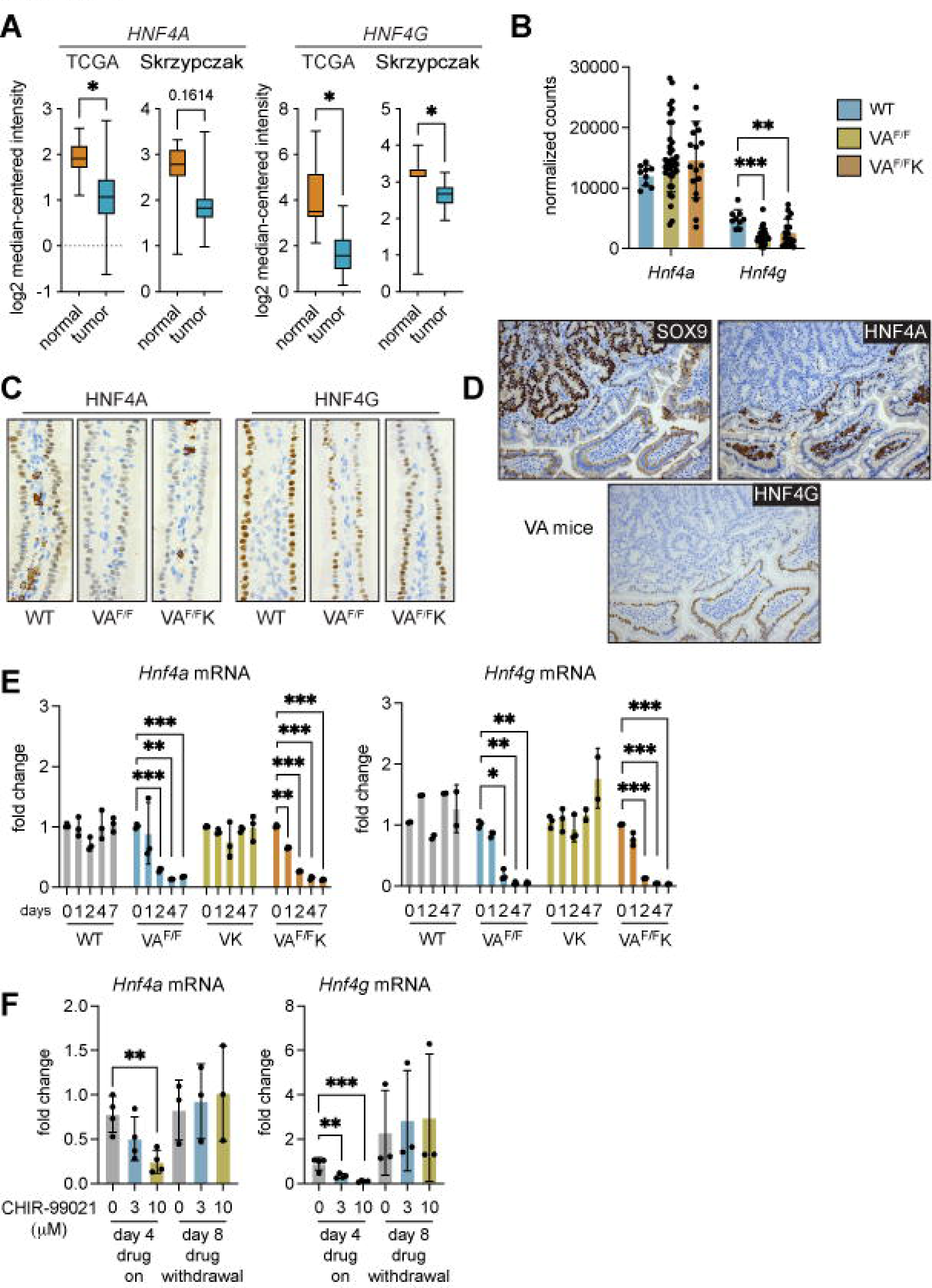
Activation of β-catenin decreases *Hnf4a and Hnf4g* expression. (A) Expression of *HNF4A* and *HNF4G* in normal human colonic tissue and tumor tissue from TCGA (n = 19 normal, 101 tumor) and Skrypczak (n = 24 normal, 45 tumor) datasets. Data presented as median ± min/max. \**p* < 0.05 as determined by Mann-Whitney U test. (B) *Hnf4a* and *Hnf4g* expression determined by RNAseq analysis of small intestine in WT, VA^F/F^ and VA^F/F^K mice. Each dot represents one mouse. Data presented as mean ± SD. \**p* < 0.05 as determined by one-way ANOVA followed by Dunnett’s posthoc test. (C) Representative images of HNF4A and HNF4G protein expression in small intestine of WT, VA^F/F^ and VA^F/F^K mice. (D) Representative images of intestinal tissue from 4 tumor-bearing VA mice stained for SOX9, HNF4A and HNF4G. (E) Fold change in expression levels of *Hnf4a* and *Hnf4g* in WT, VA^F/F^, VK and VA^F/F^K organoids. Gene expression was measured at indicated days post tamoxifen treatment. Each dot represents one organoid from one mouse. Data are presented as mean ± SD. \**p* < 0.05, \*\**p* < 0.01 and \*\*\**p* < 0.001 as determined by one-way ANOVA followed by Dunnett’s posthoc test. (F) Fold change in expression levels of *Hnf4a* and *Hnf4g* in WT organoids treated with 3 or 10 μM CHIR-99021 for indicated days. Each dot represents one organoid from one mouse. Data presented as mean ± SD. \*\**p* < 0.01, \*\*\**p* < 0.001 as determined by one-way ANOVA followed by Dunnett’s posthoc test.

We next questioned whether expression of *Hnf4a* and *Hnf4g* mRNAs were affected by WNT signaling, by examining mRNA levels in the SI of WT, VA^F/F^ and VA^F/F^K mice. This analysis showed that *Hnf4a* levels were similar between normal intestinal tissue and *Apc*-deficient tissue, whereas *Hnf4g* levels were reduced in *Apc*-deficient tissue (Figure 6B). Immunohistochemistry on these short-term models revealed that nuclear staining of both HNF4A and HNF4G was reduced or even absent from epithelial cells in the villus of VA^F/F^ and VA^F/F^K tissue when compared to WT tissue (Figure 6C). The addition of mutant KRAS to *Apc* loss had no influence over decreased expression of HNF4A and HNF4G. The discrepancy between *Hnf4a* mRNA levels and HNF4A protein levels may be explained by expression of HNF4A^+^ stromal cells in the lamina propria. These data show that expression of HNF4A and HNF4G is rapidly reduced or lost completely in cells with high β-catenin activity.

End-stage tumors from VA mice were evaluated for the presence of HNF4A and HNF4G. Nuclear SOX9 staining was used to identify WNT-high tumors. We found that HNF4A and HNF4G were completely absent from cancer cells, while normal adjacent epithelial cells maintained nuclear HNF4A and HNF4G staining (Figure 6D). This pattern of expression mimicked loss of *Btnl1* staining in tumors from the same mouse model (Figure 3A).

We used the organoid transformation assay to test the kinetics of *Hnf4a* and *Hnf4g* down-regulation after β-catenin activation. After tamoxifen treatment, expression of these molecules remained constant in WT organoids (Figure 6E). The deletion of *Apc* resulted in reduced expression of *Hnf4a* and *Hnf4g* by day 2, which was 2 days earlier than was observed for *Btnl* mRNA down-regulation, as shown in Figure 4E. Induction of oncogenic KRAS had no effect on *Hnf4a* and *Hnf4g* gene expression, but the combination of *Apc* deletion and mutant KRAS expression in VA^F/F^K organoids resulted in a down-regulation of *Hnf4a* and *Hnf4g* by day 1 or 2 (Figure 6E). These observations indicate that suppression of *Hnf4a* and *Hnf4g* RNAs by β-catenin precedes the down-regulation of *Btnl* gene expression. Treatment of WT organoids with the GSK3β inhibitor, CHIR-99021, reduced expression of *Hnf4a* and *Hnf4g* (Figure 6F). As observed with *Btnl1, Btnl2, Btnl4* and *Btnl6* expression (Figure 4F), the inhibition of *Hnf4a* and *Hnf4g* mRNA was reversible after withdrawal of CHIR-99021 with expression levels returning to normal on day 8 (Figure 6F).

Together, these data are consistent with the notion that high WNT signaling suppresses *Btnl1/2/4/6* gene expression, via down-regulation of HNF4A and HNF4G.

### The HNF4-BTNL-γδ T cell axis is restored in tumors by interference with β-catenin activity

The β-catenin transcription factor complex consists of several components including B cell lymphoma 9 (BCL9) and BCL9-like (BCL9L) (40, 41), whose deletion in mouse tumor models abrogates β-catenin-mediated transcription (26, 42). Therefore, we investigated expression levels of *Hnf4a, Hnf4g, Btnl1, Btnl2, Btnl4* and *Btnl6* in tissue where *Apc* is deleted and *Bcl9* and *Bcl9l* are absent. For this purpose, we examined RNAseq data from intestinal tissue of VA^F/F^ mice and VA^F/F^;*Bcl9^F/F^;Bcl9l^F/F^* mice that were treated with tamoxifen for 4 days to induce Cre recombinase. This analysis showed that *Hnf4g*, *Btnl1* (although not statistically significant), *Btnl2* and *Btnl4* levels are higher in VA^F/F^;*Bcl9^F/F^;Bcl9l^F/F^* intestinal tissue, while *Hnf4a* mRNA remained unchanged (Figure 7A). *Btnl6* could not be detected in this dataset.

**Figure 7.**
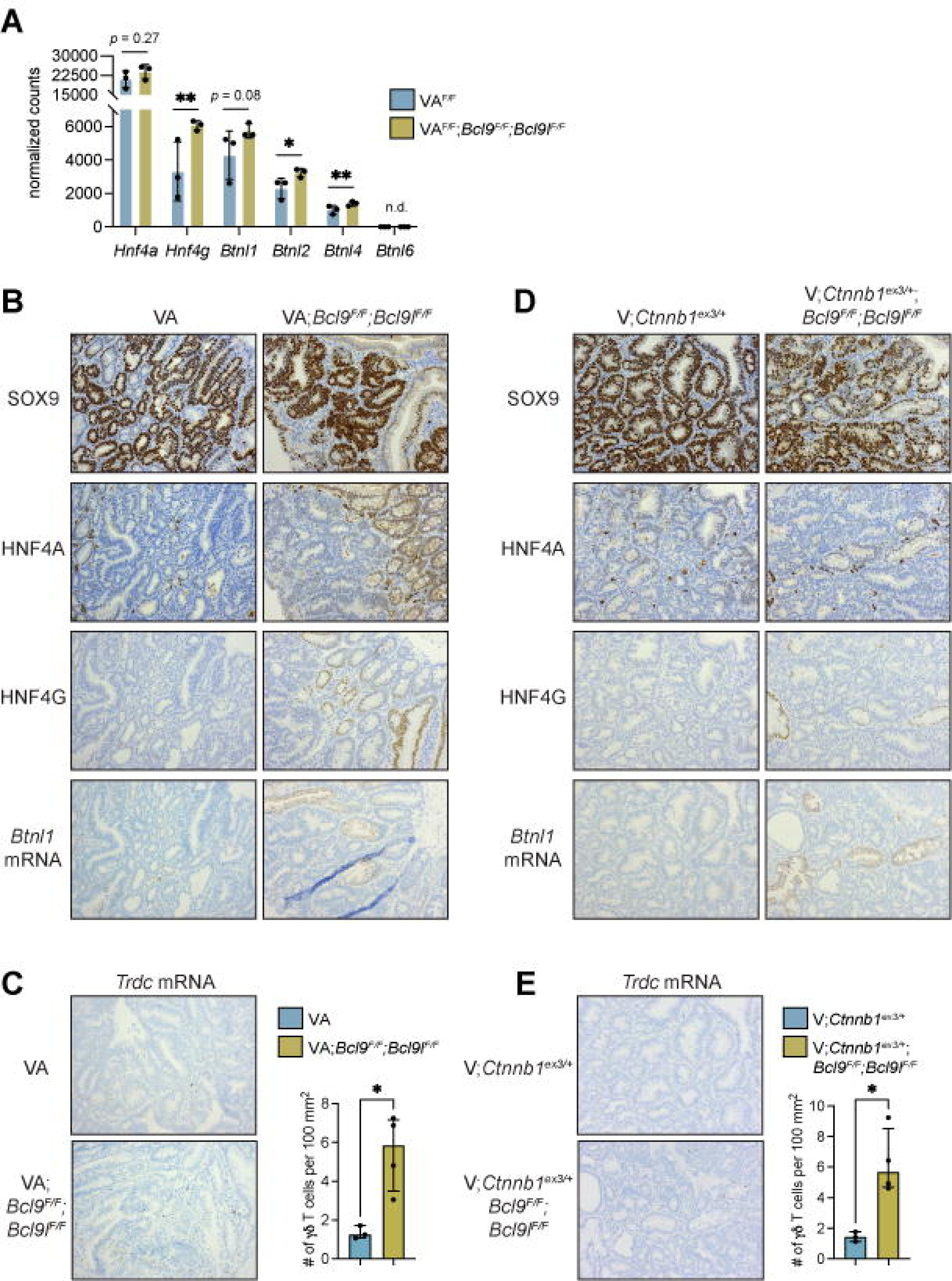
Inhibition of β-catenin transcriptional activity increases expression of HNF4A, HNF4G *and* butyrophilin-like molecules. (A) Gene expression of indicated molecules in VA^F/F^ and VA^F/F^;*Bcl9^F/F^;Bcl9l^F/F^* intestinal tissue generated from RNAseq data. Each dot represents one mouse. Data presented as mean ± SD. \**p* < 0.05, \*\**p* < 0.01 as determined by unpaired t test. n.d. = not detected. (B) Representative images of SOX9, HNF4A, HNF4G and *Btnl1* expression in tumors from 4 VA and VA;*Bcl9^F/F^;Bcl9l^F/F^*mice. (C) Representative images of *Trdc* expression in tumors from 4 VA and VA;*Bcl9^F/F^;Bcl9l^F/F^* mice. Graphic representation of γδ T cell numbers in tumors. Each dot represents one mouse. Data presented as mean ± SD from 100 mm^2^ tissue. \*\*\**p* < 0.001 as determined by unpaired t test. (D) Representative images of SOX9, HNF4A, HNF4G and *Btnl1* expression in tumors from 3 *Villin-Cre^ERT2^;Ctnnb1^ex3/+^* (V;*Ctnnb1*^ex3/+^) and V;*Ctnnb1*^ex3/+^;*Bcl9^F/F^;Bcl9l^F/F^* mice. (E) Representative images of *Trdc* expression in tumors from 3 V;*Ctnnb1*^ex3/+^ and V;*Ctnnb1*^ex3/+^;*Bcl9^F/F^;Bcl9l^F/F^*mice. Graphic representation of γδ T cell numbers in tumors. Each dot represents one mouse. Data presented as mean ± SD from 100 mm^2^ tissue. \**p* < 0.05 as determined by unpaired t test.

End-stage tumors from VA and VA;*Bcl9^F/F^;Bcl9l^F/F^*mice were assessed for expression of HNF4A, HNF4G and *Btnl1* mRNA. SOX9 was used to detect WNT-high cancer cells. Expression of HNF4A, HNF4G and *Btnl1* mRNA was absent from tumors in VA mice (Figure 7B). By contrast, nuclear expression of HNF4A and HNF4G as well as *Btnl1* mRNA was apparent in some but not all areas of tumors from VA;*Bcl9^F/F^;Bcl9l^F/F^* mice (Figure 7B). Previous reports indicate that recombination of *Bcl9^F/F^;Bcl9l^F/F^* alleles is inefficient in these mice (26), which provides an explanation for the sporadic expression pattern of HNF4A, HNF4G and *Btnl1* mRNA in these tumors. To determine whether the restoration of HNF4A, HNF4G and *Btnl1* expression in tumors from VA;*Bcl9^F/F^;Bcl9l^F/F^* mice affected tumor-infiltrating γδ T cells, we quantified these cells in tumor tissue. This analysis showed that γδ T cells are more abundant in tumors from VA;*Bcl9^F/F^;Bcl9l^F/F^* mice than VA mice (Figure 7C).

Another colon cancer model, *Villin-Cre^ERT2^;Ctnnb1*^ex3/+^ (V;*Ctnnb1*^ex3/+^) mice, was used to validate these findings, where mutant β-catenin is used to drive tumor formation (26). Nuclear SOX9 expression was used to identify WNT-high cancer cells. In this model, nuclear HNF4A expression was evident in cancer cells (albeit weak expression), while HNF4G and *Btnl1* expression were lost from tumors (Figure 7D). V;*Ctnnb1*^ex3/+^ mice were crossed with *Bcl9^F/F^;Bcl9l^F/F^* mice. Tumors from V;*Ctnnb1*^ex3/+^*;Bcl9^F/F^;Bcl9l^F/F^*mice retained expression of HNF4A. Nuclear HNF4G and *Btnl1* mRNA expression could be observed in overlapping regions of tumors (Figure 7D); although, staining was sporadic as in tumors from VA;*Bcl9^F/F^;Bcl9l^F/F^*mice (Figure 7B). We quantified tumor-infiltrating γδ T cells in these mice. Here, we found that tumor-infiltrating γδ T cells are more abundant in tumors from V;*Ctnnb1*^ex3/+^*;Bcl9^F/F^;Bcl9l^F/F^*mice than V;*Ctnnb1*^ex3/+^ mice (Figure 7E). These data indicate that inhibition of β-catenin signaling reverses HNF4A/HNF4G-driven *Btnl* gene expression and exclusion of Vγ7^+^ cells in tumors.

## DISCUSSION

γδIELs preserve normality in gut tissue, providing protection from invading pathogens and restraining proliferation of mutated epithelial cells (1-4,8). γδ T cell infiltration into colorectal tumors correlates with good prognosis and extended survival (23, 43), but their abundance decreases as disease stage progresses (24, 44). This study shows how immunosurveillance by γδIELs in the gut is disrupted by dysfunctional WNT signaling in cancer cells. We found that normal intestinal epithelial cells utilize HNF4G (most likely dimerized with HNF4A) to induce expression of *Btnl* gene expression. However, activation of β-catenin through mutations in the *Apc* tumor suppressor gene leads to suppression of HNF4 transcription factors, preventing the expression of *Btnl1/2/4/6* genes. We show that inhibition of β-catenin in tumors restores HNF4G-mediated *Btnl* gene expression and tumor-infiltrating γδ T cells. Overall, we provide a mechanism of evasion from anti-tumor immunosurveillance by unconventional T cells.

The biological basis for evasion from γδIEL immunosurveillance shown herein revolves around WNT-driven dedifferentiation of cancer cells towards a stem cell-like state. Dysregulated WNT signaling fosters the conversion of cancer cells towards a less differentiated phenotype reminiscent of LGR5^+^ stem cells that reside in intestinal crypts. LGR5^+^ stem cells, like cancer cells, fail to express HNF4G and BTNL molecules (11), making crypt regions and tumors immune privileged sites, devoid of γδIELs. Most work in the cancer dedifferentiation area has focused on immune escape from conventional CD8^+^ T cells (45, 46), with loss of differentiation-associated antigens being one mechanism of immune evasion (47, 48). Moreover, there is a strong association between the WNT pathway, dedifferentiation and CD8^+^ T cell suppression. Pan-cancer analyses of human tumor samples have shown that enrichment of the WNT pathway and its transcriptional signatures is associated with low T cell infiltration (49, 50). In mouse models of melanoma, hepatocellular carcinoma and mammary cancer, WNT signaling directly suppresses CD8^+^ T cell anti-tumor responses through various mechanisms. WNT signaling prevents dendritic cell activation via inhibition of chemokines within the tumor microenvironment (18–20), induces immunosuppressive mediators from dendritic cells to establish immune tolerance (51–53), fosters CD8^+^ T cell-suppressive neutrophils (21,54–56), as well as directly represses CD8^+^ T cells and NK cells through engagement with the LRP5 receptor (57). In addition to WNT signaling negatively affecting dendritic cells, CD8^+^ T cells and NK cells, our study offers evidence that WNT signaling can also modulate the interaction between innate-like γδ T cells and cancer cells to initiate immune escape from γδIELs.

Redifferentiation of colon cancer cells could reengage γδIEL immunosurveillance, and strategies to achieve redifferentiation could benefit γδIEL-based cancer immunotherapies. At the same time, redifferentiation would slow the proliferative signals induced by β-catenin in cancer cells. Our data suggest that redifferentiation may be possible given that inhibition of β-catenin transcriptional activity by deletion of BCL9 and BCL9L results in re-expression of HNF4G and BTNL molecules and increased numbers of γδ T cells in tumors. This notion is supported by data from other disease settings. A recent report showed that individuals with celiac disease exhibit a loss of *BTNL8* expression concomitant with a loss of Vγ4^+^Vδ1^+^ IELs, but elimination of dietary gluten could restore *BTNL8* expression (58). Together, our two studies emphasize the reversibility of *BTNL* gene expression in different disease contexts.

A question remains about how to accomplish redifferentiation for all mutational subtypes of colorectal cancer. WNT ligand-dependent tumors are susceptible to drugs that inhibit extracellular WNT ligands, such as porcupine inhibitors (59) or R-spondin blocking antibodies (60). As such, these may be used to lower β-catenin transcriptional activity, induce redifferentiation and reengage γδIELs. However, the usefulness of these drugs in WNT ligand-independent tumors is limited, and drugs specific for WNT ligand-independent tumors are scarce. Given the importance of HNF4G in driving enterocyte differentiation (33, 37) and regulating *Btnl* gene expression showed herein, drugs that increase this transcription factor and its binding partners should be prioritized in the cancer setting. It should also be noted that such strategies to restore γδIEL immunosurveillance will be anatomical site-specific. BTNL-responsive mouse Vγ7^+^ cells and human Vγ4^+^ cells are restricted to the gut, so reengagement of endogenous γδIELs will not be possible for liver metastasis. Therefore, such a strategy would be most efficacious in the primary setting, perhaps for minimal residual disease after surgery or radiotherapy.

## ACKNOWLEDGEMENTS

We are grateful to Laura Machesky (CRUK Beatson Institute), Karen Edelblum (Rutgers University) and Catherine Winchester (CRUK Beatson Institute) for advice. We thank David Bryant (University of Glasgow) for lentiviral reagents. We thank the Core Services and Advanced Technologies at the Cancer Research UK Beatson Institute, with particular thanks to the Histology Core Facility and Biological Services Unit.

This work was supported by grants from Wellcome Trust (Seed Award 208990/Z/17/Z to SBC and Senior Clinical Research Fellowship 206314/Z/17/Z to SJL); Cancer Research UK Glasgow Centre (A25142 to SBC); Marie Skłodowska Curie Actions European Fellowship (GDCOLCA 800112 to TS); Naito Foundation Grant for Research Abroad (to TS); Medical Research Council (MR/R502327/1 to SBC & JE and MR/R502327/1 to JE); Greater Glasgow and Clyde endowment (306620-01 to JE); Cancer Research UK (Early Detection Project Grant (A29834 to PD); and the NIH (R01DK121915 and R01CA190558 to MV). AH was supported by Cancer Research UK core funding at the Francis Crick Institute (FC001093). OJS and KB were supported by Cancer Research UK core funding at the Cancer Research UK Beatson Institute (A17196 and A31287).

## AUTHOR CONTRIBUTIONS

Conceptualization, TS, AK, AJ, JE, AH, OJS, SBC; Methodology, TS, AK, RR, AG, DG, EGV, HLB-D, SJL, AH, OJS, SBC; Formal Analysis, TS, AK, HH, RB, NCR, AG, LC, MV, KG, RW, AJ, NR, PDD, SBC; Investigation, TS, AK, RR, HH, RB, NCR, AG, LC, MV, DG, EGV, HLB-D, KG, AHK, CK, DA, RW, AJ, NR, SJL, PDD, JE, SBC; Resources, TS, AK, RR, LC, MV, DG, EGV, HLB-D, AHK, CK, CT, DA, SJL, JE, OJS, SBD; Data Curation, RB, LC, MV, KG, AHK, CK, CT, DA, AR, PDD, JE; Writing – Original Draft, TS, SBC; Writing – Review & Editing, TS, AK, RR, HH, RB, NCR, AG, LC, MV, DG, EGV, HLB-D, KG, AHK, CK, CT, DA, RW, AJ, NR, KB, AR, SJL, PDD, JE, AH, OJS, SBC; Visualization, TS, AK, HH, RB, NCR, AG, LC, MV, KG, SBC; Supervision, TS, AK, RR, MV, CK, KB, AR, SJL, PDD, JE, AH, OJS, SBC; Funding Acquisition, TS, SJL, PDD, JE, AH, OJS, SBC.

## METHODS

### Mice

Animal experiments were carried out in line with the Animals (Scientific Procedures) Act 1986 and the EU Directive 2010 and sanctioned by Local Ethical Review Process. All mice were maintained on the C57BL/6J background at the Cancer Research UK Beatson Institute under licence 70/8645 and PP6345023 to Karen Blyth and 70/8646 and PP3908577 to Owen Sansom, except *Vil1-Grem1* mice and *Lgr5-Cre^ERT2^;Rspo3^INV^*mice which were maintained at the Functional Genetics Facility, Wellcome Centre for Human Genetics, University of Oxford (P0B63BC4D to Simon Leedham). Mice were bred and housed in individually ventilated cages under specific pathogen-free conditions on a 12/12-hour light/dark cycle and fed and watered *ad libitum*. Both male and female mice of at least 6 weeks old and ≥20 kg were used for experiments.

The alleles used were as follows: Villin-Cre^ERT2^ (61), Apc^580S^ (62), Kras^G12D^ (63), Bcl9^F/F^, Bcl9l^F/F^ (64), Ctnnb1^ex3/+^ (65). The generation of Villin-Cre^ERT2^;Apc^F/+^ (VA) mice, Villin-Cre^ERT2^;Apc^F/+^;Kras^G12D/+^ (VAK) mice, Villin-Cre^ERT2^;Apc^F/F^ (VA^F/F^) mice, Villin-Cre^ERT2^;Apc^F/F^;Kras^G12D/+^ (VA^F/F^K) mice, VA;Bcl9^F/F^;Bcl9l^F/F^ mice, V;Ctnnb1^ex3/+^ mice, and V;Ctnnb1^ex3/+^;Bcl9^F/F^;Bcl9l^F/F^ mice has been described (25,26,66,67). Cre negative mice were used as controls. Recombination in these tumor models was induced by a single intraperitoneal injection of 80 mg/kg tamoxifen. Mice were aged until they showed clinical signs (anemia, hunching and/or weight loss). Tumors were scored macroscopically after fixation of opened intestinal tissue. Tumor burden was calculated by summing the area of all tumors. Recombination of VA^F/F^ and VA^F/F^K short-term models was induced using intraperitoneal injections of 80 mg/kg tamoxifen for 2 consecutive days; wild-type control mice received the same dosing regimen. Mice were sacrificed 3 or 4 days after the first injection. Generation of Lgr5-Cre^ERT2^;Rspo3^INV^ mice has been described (39). Recombination in this model was induced by intraperitoneal injection of 1 mg tamoxifen for 5 consecutive days. Mice were aged until they showed clinical signs (anemia, hunching and/or weight loss). The porcupine inhibitor LGK-974 was administered by daily oral gavage at 1 mg/kg in 0.5% hydroxypropyl methylcellulose. Vil1-Grem1 mice and Btnl1^—/—^ mice were generated as described previously (11, 38).

### Immunohistochemistry and *in situ* hybridization

Tissues were fixed overnight in 10% neutral buffered formalin, then embedded in paraffin. Staining was performed on 4 µm sections, which had been heated at 60 ⁰C for 2 hours. Primary antibodies used for IHC were as follows: CDX1 (1:250; Invitrogen #PA5-23056), CDX2 (1:200; Abcam #ab76541), HNF4A (1:10,000; Perseus Proteomics #pph1414-00), HNF4G (1:1000; Novus Biologicals #NBP1-82531), SOX9 (1:500; Millipore #AB5535). HNF4A and SOX9 were detected by an Agilent AutostainerLink48 using high pH citrate buffer (Target Retrieval Solution, Aglient #K8004/K8005) and peroxidase blocking. CDX1, CDX2, and HNF4G were detected on a Leica Bond Rx autostainer, using ER2 antigen retrieval solution (Leica #AR9640). For RNAscope, the following probes were used from Advanced Cell Diagnostics: *Btnl1* (436648) and *Trdc* (449358). Staining was performed on a Leica Bond Rx autostainer according to Advanced Cell Diagnostics instructions. Images were acquired with an Olympus BX51 or Zeiss Axio Imager.A2 microscope. For each antibody or RNAscope probe, staining was performed on tissue sections from at least three mice of each genotype, and representative images are shown for each staining. The average number of γδ T cells was determined by HALO image analysis software (Indica Labs) in 10^6^ μm^2^ tissue from 1-5 villi or tumors within each mouse.

### Gene expression analysis of mouse tissue

RNA-seq data from wild-type, VA^F/F^, and VA^F/F^K mouse intestinal tissue was generated for previous studies (ArrayExpress E-MTAB-7546) (25,26,68). Analysis of these data was performed as previously described where raw counts per gene were determined using FeatureCounts version 1.6.4 (68). Differential expression analysis was performed using the R package DESeq2 version 1.22.2, and principal component analysis was performed using R base functions. RNA-seq data from wild-type, *Hnf4a*^Δ*/*Δ^, *Hnf4g*^Δ*/*Δ^ and *Hnf4a*^Δ*/*Δ^*;Hnf4g*^Δ*/*Δ^ mice was analyzed as previously described (33); these data are available (GSE112946).

### Flow cytometry

Tumors and 1 cm^2^ of jejunum were cut into small pieces using the McIlwain™ Tissue Chopper and digested on the gentleMACS™ Octo Dissociator with Heaters (program, 37C_m_TDK_1) using the mouse Tumor Dissociation Kit (Miltenyi Biotec) according to the manufacturer’s instructions, and prepared cells were resuspended in 0.5% BSA in PBS. Cells were stained in the brilliant stain buffer (BD Biosciences) containing antibodies for 30 min at 4 °C in the dark. The following antibodies were used:

| Antigen | Clone | Conjugate | Source | Catalogue | Dilution |
| --- | --- | --- | --- | --- | --- |
| CD19 | 1D3 | APC-eFluor780 | eBioscience | 47-0193-82 | 1:400 |
| CD3ε | 17A2 | BV650 | BioLegend | 100229 | 1:100 |
| CD8α | 53-6.7 | BUV805 | BD | 564920 | 1:50 |
| EpCAM | G8.8 | APC-eFluor780 | eBioscience | 47-5791-82 | 1:100 |
| TCR delta | GL3 | FITC | eBioscience | 11-5711-85 | 1:200 |

Dead cells were identified with Zombie NIR Fixable Viability dye (Biolegend). Cells were acquired using a 5-laser BD LSRFortessa flow cytometer with DIVA software (BD Biosciences). Data were analyzed using FlowJo Software version 9.9.6.

### Human patient cohorts and immunohistochemistry

The Scotland cohort was assembled from 1030 patients who had undergone a resection for Stage I-IV colon cancer between 1997 and 2007 at the Glasgow Royal Infirmary, Western Infirmary or Stobhill Hospital in Glasgow, UK. Tumors were staged using the 5^th^ edition of AJCC/UICC-TNM staging system. A sub-cohort of 144 samples were selected for IHC, and tissue was available from 142 patients where both tumor and normal adjacent tissue was visible. The Norway cohort was assembled from 299 patients who had undergone a resection for Stage II-III colon cancer between 2000 and 2020 at the Southern Hospital Trust in Norway. Tumors were staged using the 5^th^ edition of AJCC/UICC-TNM staging system from 2000 to 2009, the 7^th^ edition from 2010 to 2017, and the 8^th^ edition thereafter. A sub-cohort of 84 samples were selected for IHC, and tissue was available from 71 patients where both tumor and normal adjacent tissue was visible. The Thailand cohort was assembled from 411 patients who had undergone a resection for Stage I-IV colon cancer between 2009 and 2016 at hospitals in Thailand. These samples were approved by the Siriraj Institution Review Board (COA no.Si544/2015). Tumors were staged using the 6^th^ or 7^th^ editions of AJCC/UICC-TNM staging system. A sub-cohort of 136 samples were selected for IHC, and tissue was available from 122 patients where both tumor and normal adjacent tissue was visible. Across all cohorts, patients were excluded if they had received neoadjuvant chemotherapy or died within 30 days of surgery. The cohorts consisted of the following clinicopathological characteristics:

|  | *Scotland* | *Norway* | *Thailand* |
| --- | --- | --- | --- |
| **Sex** |  |  |  |
| Female | 73 (51%) | 41 (49%) | 60 (44%) |
| Male | 71 (49%) | 43 (51%) | 76 (56%) |
| **Age** |  |  |  |
| >=65 | 90 (62%) | 68 (81%) | 69 (51%) |
| <65 | 54 (38%) | 16 (19%) | 67 (49%) |
| **Tumor site** |  |  |  |
| Right | 67 (47%) | 48 (57%) | n.a. |
| Left | 46 (32%) | 36 (43%) |  |
| Rectum | 31 (22%) | 0 (0%) |  |
| **TMN Stage** |  |  |  |
| I | 16 (11%) | 0 (0%) | 0 (0%) |
| II | 60 (42%) | 51 (61%) | 24 (18%) |
| III | 63 (44%) | 33 (39%) | 61 (45%) |
| IV | 5 (3%) | 0 (0%) | 37 (27%) |
| **Differentiation** |  |  |  |
| Moderate/Well | 131 (91%) | 65 (77%) | 133 (98%) |
| Poor | 13 (9%) | 16 (19%) | 0 (0%) |
| Missing | 0 (0%) | 3 (3.6%) | 3 (2.2%) |
| **MMR Status** |  |  |  |
| Proficient | 116 (81%) | n.a. | n.a. |
| Deficient | 25 (17%) |  |  |
| Missing | 3 (2.1%) |  |  |

IHC was performed on full tissue sections with citrate buffer (pH 6.0) antigen retrieval with standard protocols, using an anti-TCRδ antibody (1:300; clone H-41, Santa Cruz #sc-100289, lot K1318 or K2618) previously validated (69). Scoring of γδ T cells was conducted using VisioPharm® software. The first level of tissue compartments (primary tumor, adjacent normal tissue) was manually annotated. A tissue classifier was built using RGB and haematoxylin features with the application of a K-means clustering algorithm and was trained using sections from all cohorts. A pan-lymphocyte detector was built using a five-pixel mean filter applied to the chromogenic DAB feature and a dual haematoxylin feature consisting of a polynomial smoothing filter and a polynomial Laplace filter at a field size of 15 pixels at an order of two. The output metric is defined as the % of total cells within an analysed region that are positively identified as the target cell type.

### Gene expression analysis in human tumors

TempO-Seq (Biospyder Technologies, Carlsbad, CA, USA) whole transcriptome profiling was performed on 82 patients from the Scotland cohort, according to the manufacturer’s instructions using whole FFPE tissue sections. 77 out of the 82 had matched γδ T cell IHC data. FFPE tissue were deparaffinised prior to tissue digestion. Crude tissue lysates were used as input for whole transcriptome analysis using the Human Whole Transcriptome v2.0 panel. Detector oligos, consisting of a sequence complementary to an mRNA target plus a universal primer landing site, were annealed in immediate juxtaposition to each other on the targeted RNA template and ligated (70). Amplification of ligated oligos were performed using a unique primer set for each sample, introducing a sample-specific barcode and Illumina adaptors. Barcoded samples were pooled into a single library and run on an Illumina HiSeq 2500 High Output v4 flowcell. Sequencing reads were demultiplexed using BCL2FASTQ software (Illumina, USA). FASTQ files were aligned to the Human Whole Transcriptome v2.0 panel, which consist of 22,537 probes, using STAR (71). Up to two mismatches were allowed in the 50-nucleotide sequencing read. Deseq2 was used to normalize raw read counts. Linear regression analysis on paired samples was performed using Prism software (version 9.3.1).

Oncomine was used to query gene expression levels of *BTNL3, BTNL8, HNF4A, HNF4G, CDX1* and *CDX2* in normal and tumor tissue from the TCGA (15) and Skrzypczak (27) cohorts. Expression levels are presented as log_2_ median-centered intensity.

The Marisa cohort consists of fresh-frozen primary tumor samples from patients with colon cancer collected and transcriptionally profiled as described previously (30). The normalized, batch corrected microarray data for the Marisa cohort were downloaded from Gene Expression Omnibus (GEO) using the accession number GSE39582. This dataset had been processed using the Robust Multi-Array Analysis (RMA) method and corrected for technical batch effects using ComBat as described previously (30). Probesets were collapsed to the gene level by selecting the probeset with the highest mean expression value across all samples for each gene using the collapseRows function (method=“MaxMean”) from the WGCNA package (72) using R (v3.3.2). Only tumor samples from patients with Stage II or III disease who did not receive adjuvant chemotherapy that had relapse-free survival data (n = 258) were used analysis.

### Organoid culture and treatment

Small intestine was harvested from mice of indicated genotypes. Organoids were generated as previously described (25, 68), cultured in Matrigel with ENR medium (Advanced DMEM/F12 containing 2 mM Glutamine, 10 mM HEPES, 1× N2 supplement, 1× B27 supplement, 50 ng/mL EGF (R&D Systems), 100 ng/mL Noggin (Peprotech), 1000 ng/mL R-spondin 1 (Peprotech), 100 U/mL of penicillin and 100 U/mL of streptomycin). Organoids were split every 2-3 days. Where indicated, organoids from wild-type (Cre negative mice), VA^F/F^, VK, and VA^F/F^K mice were treated with 1 μM 4-Hydroxytamoxifen (4-OHT, Sigma) in ENR medium for 48 hours. Organoids from wild-type mice were treated with 3 μM and 10 μM CHIR-99021 (Sigma) or DMSO as a control (1:300 dilution) in ENR medium for 6 days. Medium containing CHIR-99021 was changed every day. After 6 days, organoids were cultured in ENR medium without CHIR-99021 or DMSO for 2 days. Organoids from wild-type mice were treated with 100 μM BI-6015 (Cayman) or DMSO as a control (1:100 dilution) in ENR medium for 3 days. Cells were collected for downstream analysis on indicated day after treatment. Biological replicates were generated from individual mouse organoid lines.

Short hairpin (sh)RNA target sequences designed against *Cdx1, Cdx2, Hnf4a, Hnf4g* and *Sox9* were selected from Merck Mission shRNAs (https://www.sigmaaldrich.com/GB/en/product/sigma/shclnd). 5 sequences per gene were subcloned into the pLKO.1-Puro lentiviral backbone (https://www.addgene.org/8453/), and inserts sequenced before use. Viral supernatants were prepared following transient transfection of 293FT cells with pLKO.1 encoding shRNAs, pSPAX2 packaging vector and pVSVG envelope vector using Lipofectamine 2000 (Thermo Fisher Scientific, Waltham, MA, USA) as described (73). Two 24-hour supernatants were collected sequentially over a 48 hour period, pooled and filtered through a 0.45 μm syringe filter and then concentrated using the Lenti-X Concentrator solution (Clontech/Takara, Saint-Germain-en-Laye, France). Intestinal WT organoids were expanded 3 days prior to infection in normal growth medium supplemented with 1 μg/ml R-Spondin, 3 μM CHIR-99021 (GSK3β inhibitor), 10 μM Y27632 (ROCK inhibitor), and 1 μM Jagged-1 (Notch Ligand 1) to enrich stem and progenitor cells (74). VA^F/F^ organoids that received *Sox9* shRNAs were similarly expanded but no supplements were added. Organoids were reseeded into the same medium 24 hours before infection. Freshly concentrated viral supernatants were added directly to harvested, manually disrupted organoids in the presence of 8 μg/ml polybrene and mixtures seeded into 12 well plates coated with a fine film of Matrigel. Organoid fragments were left to attach overnight and then drained before overlaying with a fine film of Matrigel. Organoids were expanded in culture medium as above, supplemented with 1 μg/ml R-Spondin (WT organoids only) and 3 μg/ml puromycin.

### Quantitative RT-PCR

RNA was isolated from fresh intestinal organoids using Qiagen’s RNeasy kit (Manchester, UK) with on-column DNA digestion. RNA concentration and purity (cutoff = 2.0-2.2 260/280 ratio) was determined using a Thermo Scientific NanoDrop spectrophotometer with NanoDrop 2000 software. cDNA was prepared from 0.5-1 μg RNA using a Quantitect Reverse Transcription Kit (Qiagen) and diluted to 2.5 ng/mL in DEPC-treated water. For quantitative RT-PCR, 12.5 ng aliquots of cDNA were amplified in triplicate on an ABI 7500 real-time PCR machine using SyGreen Blue Mix Lo-ROX PCR master mix (PCR Biosystems, London, UK) and primers (below), all at 2.0 μM except for *Btnl1* (1 μM Fwd; 4 μM Rv), with endogenous controls *Hprt* (Mm_Hprt_1_SG; Quantitect) and β-actin (Mm_Actn_1_SG; Quantitect). Relative expression was calculated by the ΔCt method after averaging endogenous controls. Data are displayed as fold change (2^-ΔΔCt^). The following primer sequences were used for each gene: *Btnl1* forward 5’- CCGGGAACACGCTACTGTC-3’, reverse 5’-CAAACCAGGGCTACTTTCCAT-3’; *Btnl2* forward 5’-TTTGCTATGGATGACCCTGC-3’, reverse 5’-TCCTGATTGCTGCTGTGTGT-3’; *Btnl4* forward 5’-CATTCTCCTCAGAGACCCACACTA-3’, reverse 5’-GAGAGGCCTGAGGGAAGAA-3’; *Btnl6* forward 5’-CGTGTGGAGGATAATAAGGCAGA-3’, reverse 5’-TCCTTGCGCCAATCTGCATAC-3’. The other primers were purchased from QIAGEN (Quantitect Primer): *Hprt* (QT00166768); *Axin2* (QT00126539); *Lgr5* (QT00123193); *Sox9* (QT00163765); *Cdx1* (QT00265139): *Cdx2* (QT00116739); *Cd44* (QT00173404); *Hnf4a* (QT00144739); *Hnf4g* (QT00169799).

### Gene promoter analysis

Promoter sequences for mouse *Btnl1*, *Btnl2*, *Btnl4* and *Btnl6* and human *BTNL3* and *BTNL8* (12 kb upstream of ATG start site) were extracted from the UCSC Genome Browser (75). These sequences were analyzed by The Open Regulatory Annotation database (ORegAnno) (76) for putative transcription factor binding sites using ApE software.

### ChIP-seq

ChIP-seq data were generated as previously described (33), using anti-HNF4A (6 μg, Santa Cruz #sc-6556 X, lot B1015) and anti-HNF4G (6 μg, Santa Cruz #sc-6558 X, lot F0310) antibodies. ChIP-seq were visualized using IGV (77). ChIP-seq data are available GSE112946.

### Statistical analysis

An unpaired t test or the non-parametric Mann-Whitney test was used to compare two groups. One-way ANOVA was used to compare groups of three or more followed by Tukey’s or Dunnett’s posthoc test. The log-rank (Mantel–Cox) test was used to analyze Kaplan–Meier survival curves. Correlation between genes was determined using the Pearson correlation coefficient. *P* values less than 0.05 were considered statistically significant. Graphs were generated and statistical significance calculated using Prism software (version 9.3.1). The statistical tests used are indicated in figure legends. For all animal and organoid experiments, each data point represents an individual mouse or individual organoid line.

---

**Supplemental Figure 1.**
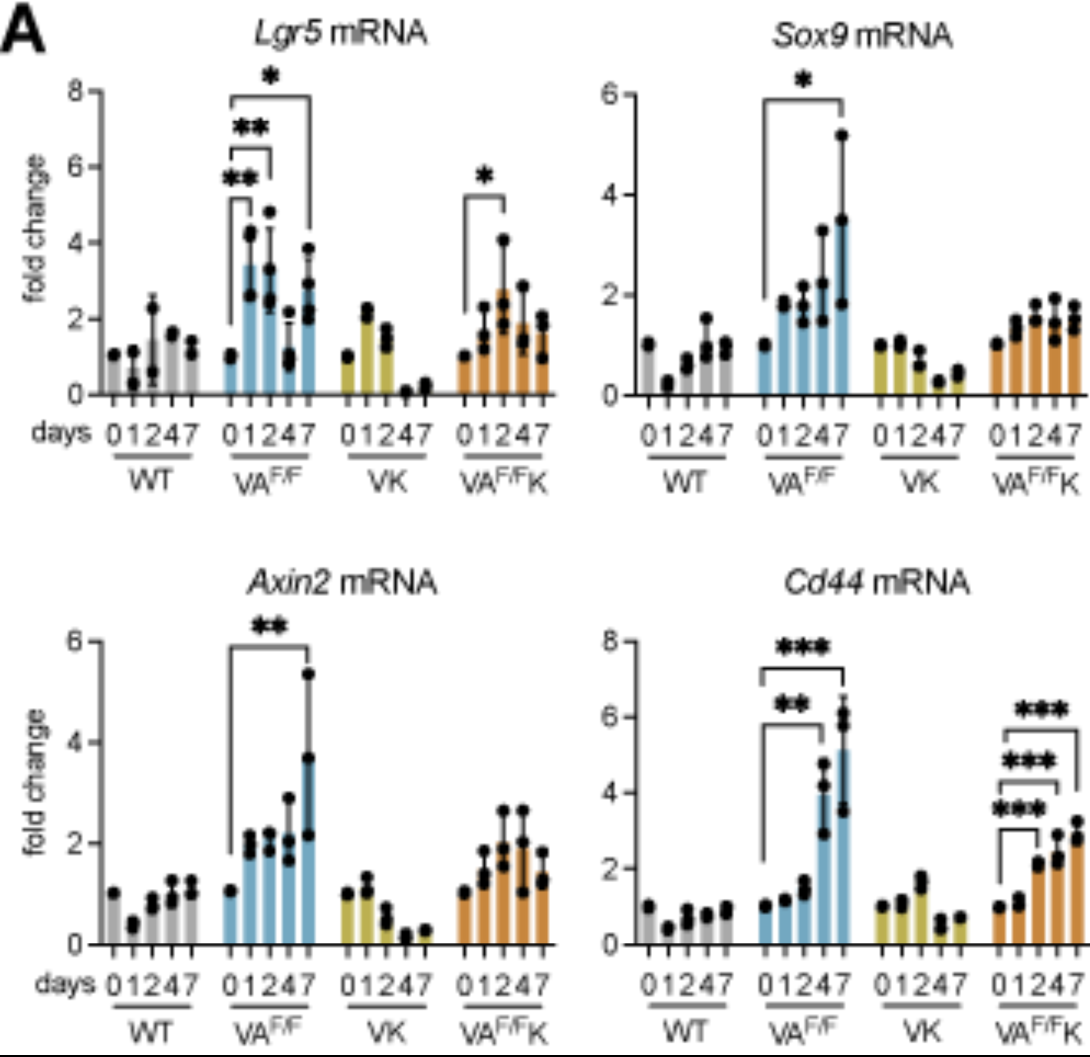
Deletion of *Apc* in organoids increases WNT target genes. (A) Fold change in expression levels of indicated genes in WT, VA^F/F^, VK and VA^F/F^K organoids. Gene expression was measured at indicated days post tamoxifen treatment. Each dot represents one organoid from one mouse. Data presented as mean ± SD. \**p* < 0.05, \*\**p* < 0.01 and \*\*\**p* < 0.001 as determined by one-way ANOVA followed by Dunnett’s posthoc test.

**Supplemental Figure 2.**
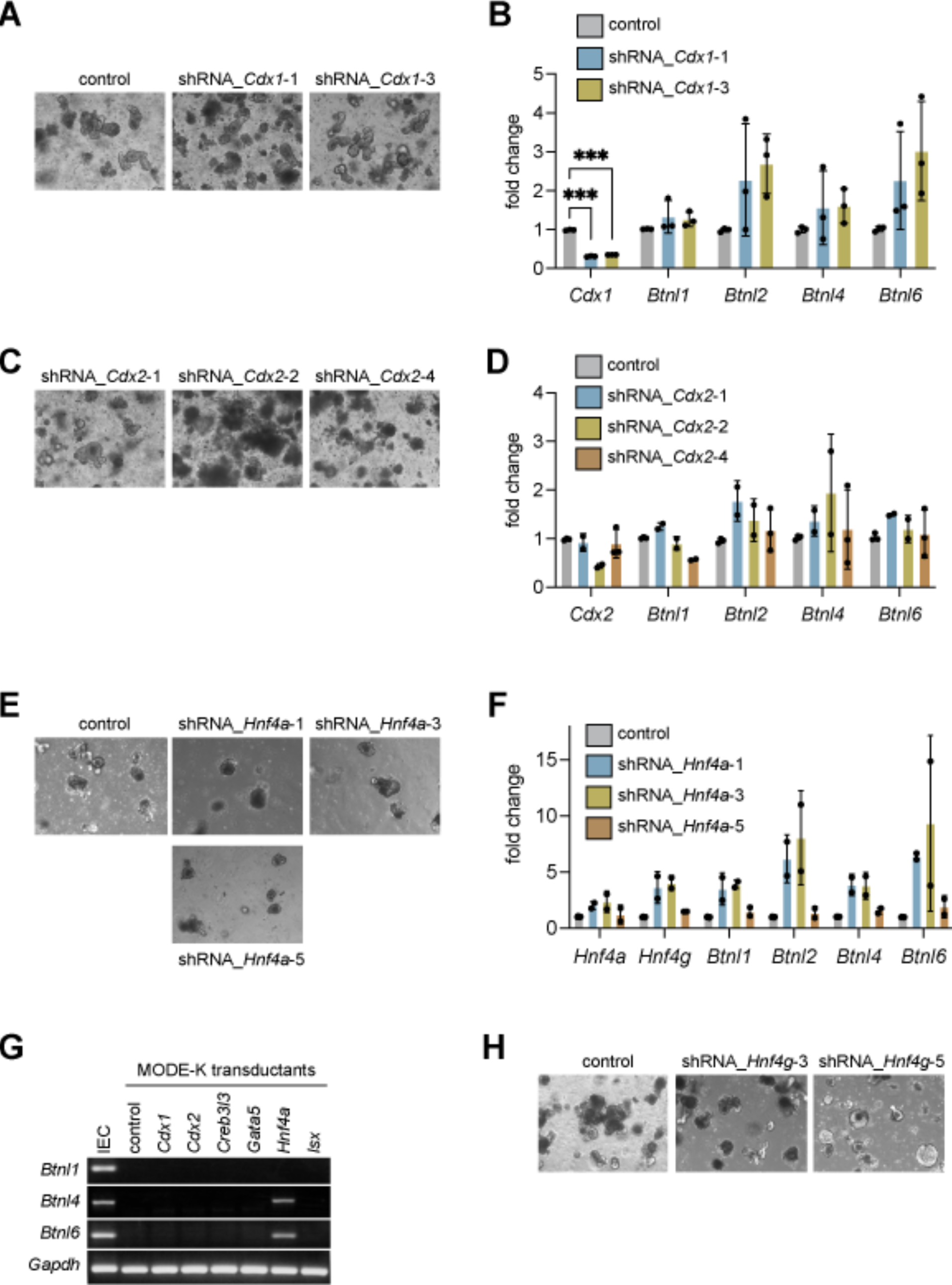
Knockdown of *Cdx1, Cdx2* and *Hnf4a* fails to influence organoid morphology or expression of *Btnl* genes. (A) Representative images of organoids from WT mice transduced with shRNA constructs against *Cdx1*. (B) Fold change in expression levels of indicated genes in WT organoids transduced with shRNA constructs targeting *Cdx1* transcripts. Each dot represents one organoid from one mouse (n = 3). Data presented as mean ± SD. \*\*\**p* < 0.001 as determined by one-way ANOVA followed by Tukey’s posthoc test. (C) Representative images of organoids from WT mice transduced with shRNA constructs against *Cdx2*. (D) Fold change in expression levels of indicated genes in WT organoids transduced with shRNA constructs targeting *Cdx2* transcripts. Each dot represents one organoid from one mouse (n = 2-3). Data presented as mean ± SD. (E) Representative images of organoids from WT mice transduced with shRNA constructs against *Hnf4a*. (F) Fold change in expression levels of indicated genes in WT organoids transduced with shRNA constructs targeting *Hnf4a* transcripts. Each dot represents one organoid from one mouse (n = 2). Data presented as mean ± SD. (G) Representative RT-PCR product bands for MODE-K cells transduced with transcription factor over-expression constructs. Intestinal epithelial cells (IEC) served as positive control, while empty vector (control) served as negative control. (H) Representative images of organoids from WT mice transduced with shRNA constructs against *Hnf4g*.

**Supplemental Figure 3.**
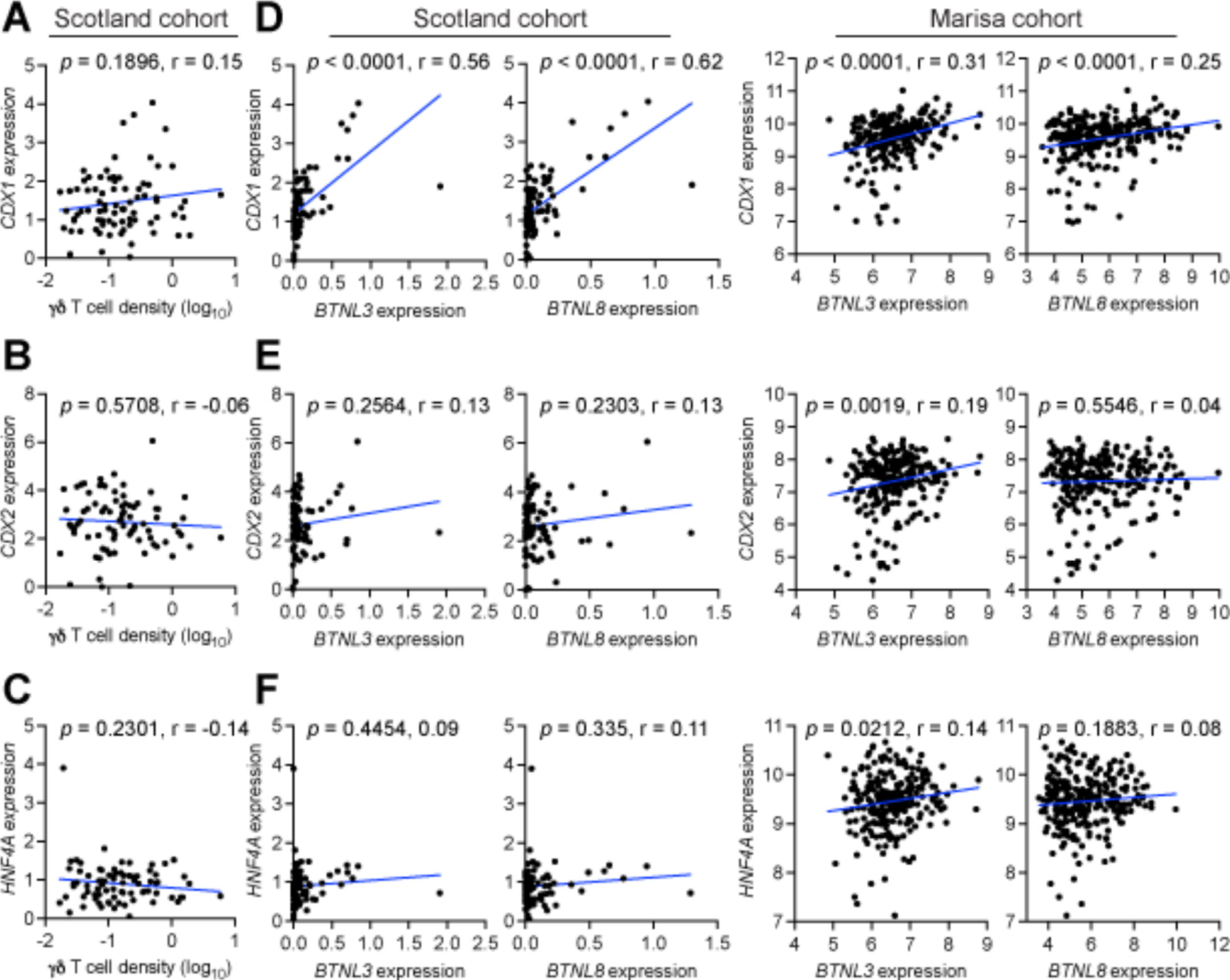
Correlation between gut-specific transcription factors, γδ T cell density and *BTNL* genes in human tumors. (A) Correlation between *CDX1* expression as determined by TempO-Seq and γδ T cell density determined by IHC in the Scotland cohort. Units on axes are normalized counts x 10^3^. Each dot represents one tumor (n = 77). *P* value and r value determined by Pearson’s correlation. (B) Correlation between *BTNL3* or *BTNL8* expression and *CDX1* expression. Units on axes are normalized counts x 10^3^. Each dot represents one tumor (n = 82 Scotland cohort, 258 Marisa cohort). *P* value and r value determined by Pearson’s correlation. (C) Correlation between *CDX2* expression as determined by TempO-Seq and γδ T cell density determined by IHC in the Scotland cohort. Units on axes are normalized counts x 10^3^. Each dot represents one tumor (n = 77). *P* value and r value determined by Pearson’s correlation. (D) Correlation between *BTNL3* or *BTNL8* expression and *CDX2* expression. Units on axes are normalized counts x 10^3^. Each dot represents one tumor (n = 82 Scotland cohort, 258 Marisa cohort). *P* value and r value determined by Pearson’s correlation. (E) Correlation between *HNF4A* expression as determined by TempO-Seq and γδ T cell density determined by IHC in the Scotland cohort. Units on axes are normalized counts x 10^3^. Each dot represents one tumor (n = 77). *P* value and r value determined by Pearson’s correlation. (F) Correlation between *BTNL3* or *BTNL8* expression and *HNF4A* expression. Units on axes are normalized counts x 10^3^. Each dot represents one tumor (n = 82 Scotland cohort, 258 Marisa cohort). *P* value and r value determined by Pearson’s correlation.

**Supplemental Figure 4.**
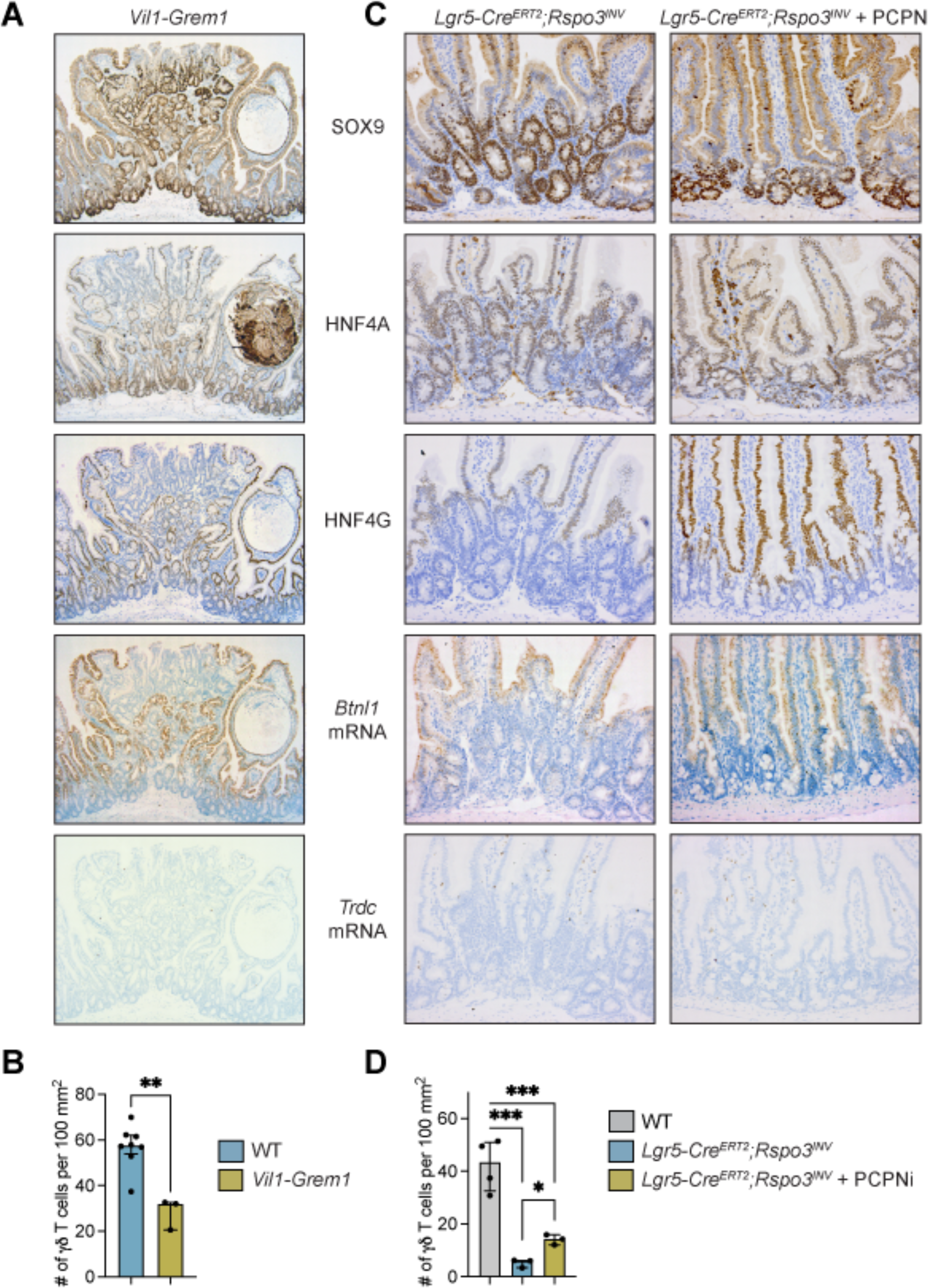
Disruption of WNT gradient in normal intestinal villi reduces γδ T cells. (A) Representative images of SOX9, HNF4A, HNF4G, *Btnl1* and *Trdc* expression in small intestine from 3 *Vil1-Grem1* mice. (B) Graphic representation of γδ T cell numbers in intestinal tissue of WT and *Vil1-Grem1* mice. Each dot represents one mouse (n = 8 WT, 3 *Vil1-Grem1*). Data presented as mean ± SD per 100 mm^2^. \*\**p* < 0.01 as determined by unpaired t test. (C) Representative images of SOX9, HNF4A, HNF4G, *Btnl1* and *Trdc* expression in small intestine from *Lgr5-Cre^ERT2^;Rspo3^INV^* mice treated with vehicle control or LGK-974 (PCPNi). (D) Graphic representation of γδ T cell numbers in intestinal tissue of WT, *Lgr5-Cre^ERT2^;Rspo3^INV^* mice and PCPNi-treated *Lgr5-Cre^ERT2^;Rspo3^INV^* mice. Each dot represents one mouse (n = 4 WT, 3 *Lgr5-Cre^ERT2^;Rspo3^INV^*, 3 *Lgr5-Cre^ERT2^;Rspo3^INV^*+ PCPNi). Data presented as mean ± SD per 100 mm^2^. \**p* < 0.05, \*\*\**p* < 0.001 as determined by one-way ANOVA followed by Tukey’s posthoc test.

