## Supplementary figures and images for "β-catenin obstructs γδ T cell immunosurveillance in colon cancer through loss of BTNL expression"

### Supplemental Figure 1

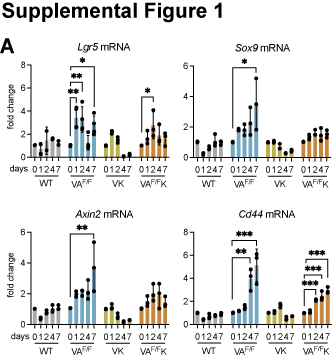

### Supplemental Figure 2

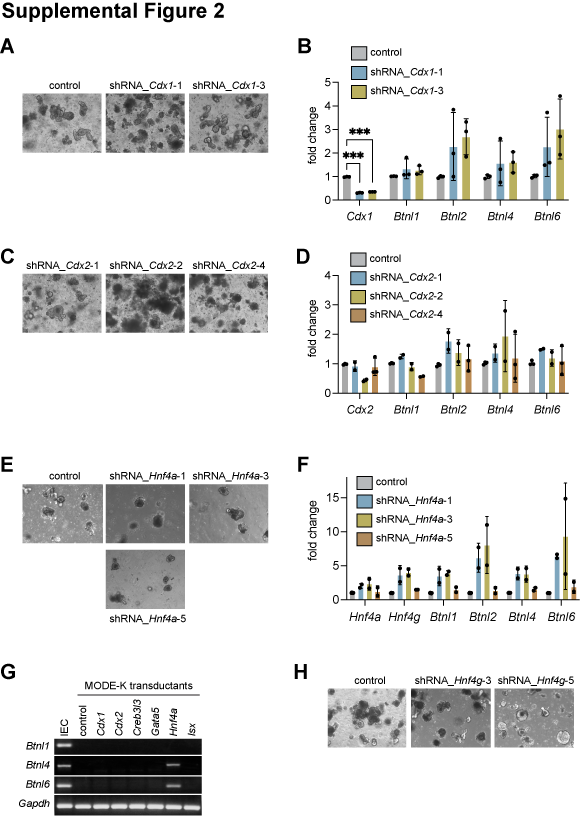

### Supplemental Figure 3

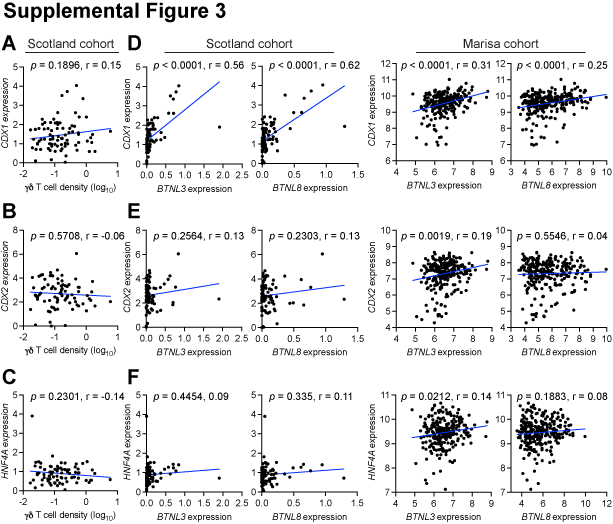

### Supplemental Figure 4

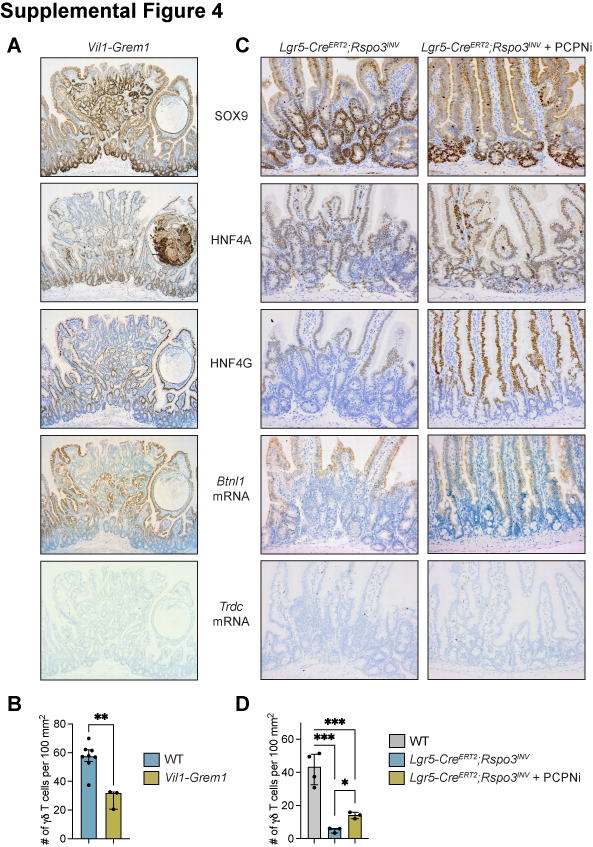
